# Chaotic aging: Intrinsically disordered proteins in aging-related processes

**DOI:** 10.1101/2023.04.22.537928

**Authors:** Vladimir D. Manyilov, Nikolay S. Ilyinsky, Semen V. Nesterov, Baraa M.G.A. Saqr, Guy W. Dayhoff, Egor V. Zinovev, Simon S. Matrenok, Alexander V. Fonin, Irina M. Kuznetsova, Konstantin K. Turoverov, Valentin Ivanovich, Vladimir N. Uversky

## Abstract

The development of aging is associated with the disruption of key cellular processes manifested as well-established hallmarks of aging. Intrinsically disordered proteins (IDPs) and intrinsically disordered regions (IDRs) have no stable tertiary structure that provide them a power to be configurable hubs in signaling cascades and regulate many processes, potentially including those related to aging. There is a need to clarify the roles of IDPs/IDRs in aging. The dataset of 1624 aging-related proteins was collected from established aging databases and experimental studies. There is a noticeable presence of IDPs/IDRs, accounting for about 36% of the aging-related dataset, which is comparable to the disorder content of the whole human proteome (about 40%). A Gene Ontology analysis of the our Aging proteome reveals an abundance of IDPs/IDRs in one-third of aging-associated processes, especially in genome regulation. Signaling pathways associated with aging also contain IDPs/IDRs on different hierarchical levels. Protein-protein interaction network analysis showed that IDPs present in different clusters associated with different aging hallmarks. Protein cluster with IDPs enrichment and high liquid-liquid phase separation (LLPS) probability has “nuclear” localization and DNA-associated functions, related to aging hallmarks: genomic instability, telomere attrition, epigenetic alterations, stem cells exhaustion. Some IDPs related to aging with high LLPS propensity were identified as “dangerous” based on the prediction of their propensity to aggregation. Overall, our analyses indicate that IDPs/IDRs play significant roles in aging-associated processes, particularly in the regulation of DNA functioning. IDP aggregation, which can lead to loss-of-function and toxicity, could be critically harmful to the cell. A structure-based analysis of aging and the identification of proteins that are particularly susceptible to disturbances can enhance our understanding of the molecular mechanisms of aging and open up new avenues for slowing it down.

## 2. Introduction

Aging is a process, occurring as a result of different uncompensated abnormities on the intracellular, inter-cellular, organ/tissue, and organismal levels [1, 2]. Since aging causes the emergence of age-related diseases, an understanding of the aging process could be a synergistic tool for combating all age-related disorders [3, 4]. Aging inside the cell is characterized by dysfunctional regulation, structure disturbances, genome instability, epigenetic changes, loss of proteostasis, mitochondrial dysfunction, and disruption of cellular metabolism [1, 2].

Intrinsically disordered proteins (IDPs) are the key proteins in signaling and stress-response processes [5] intersecting with hallmarks of aging. Indeed, phosphorylation sites [6, 7], as well as sites of many other enzymatically catalyzed posttranslational modifications (PTMs) are usually located within intrinsically disordered regions (IDRs) [8–14]. A lack of fixed structure allows IDPs/IDRs to have high binding specificity coupled with low affinity [15–17]. Consequently, signals to be ‘switched on’ and ‘switched off’ quickly. Moreover, when IDPs bind to other molecule, a portion of the binding energy is often spent to fold the IDP from the initially disordered state. This gives rise to interactions that can be weak but specific [18]. Ultimately, the structural plasticity and conformational heterogeneity of IDPs allow them to interact with a large number of often unrelated partners, placing them at the centers of protein-protein interactions networks [19–21].

It is important to find out how structural intrinsic disorder is represented in proteins associated with aging and in proteins associated with those cellular processes in which it is most pronounced. Of the aging hallmarks, “loss of proteostasis” serves as an exemplar with respect to the role IDPs play in the aging process. Aging-related loss of proteostasis is characterized by the formation of toxic protein aggregates [22]. Misfolding and aggregation of IDPs are often associated with neurodegenerative diseases [23] such as Alzheimer’s disease[24], frontotemporal lobar degeneration, and amyotrophic lateral sclerosis [25]. Additionally, disorder in IDPs/IDRs is strongly linked to other age-related diseases. Among them are cancer [26], amyloidosis [27], cardiovascular diseases [28], and diabetes [29]. Although aging itself is not a disease, it shares molecular and cellular mechanisms with age-related diseases [4].

Liquid-liquid phase separation is a common organizing principle of intracellular space and biomembranes providing dynamic adaptive responses [30]. So, maintaining the stability of stress granules and other proteinaceous membrane-less organelles (PMLOs) plays a special role in aging [31]. The proteomes of membrane-less organelles contain multiple IDPs; in addition, the disorder-based interactions among these proteins are considered to be the main driving force for the formation of PMLOs [32–43]. Protein propensity for liquid-liquid phase separation (LLPS) and aggregation could be toxic or protective [44]. On one hand, it has been shown that controlled protein aggregation has cytoprotective functions, vital for the maintenance of cell integrity and survival under adverse stress [45]. On the other hand, LLPS has protective role against protein aggregation (like in the case of stress granules) [46, 47]. LLPS and the stability of membrane-less organelles is tightly regulated in the cell, and when physicochemical conditions change, a reverse phase transition can occur. In this case, the uncontrolled formation of a gel or solid fibrous aggregates can be observed [48]. Deviation from phase transitions of the “liquid-liquid” type, which inevitably occurs with aging, can accelerate the aging of the body and the development of age-related diseases through several pathways. Cellular aging is the cause of disruption in the formation of canonical and non-canonical stress granules. In addition, in old cells, adaptation to chronic and acute stress, as well as the regulation of stress organelles, is also impaired [31].

Considering the role of intrinsic disorder in age-related diseases and accelerated aging hallmarks, it can be hypothesized that IDPs should play key roles in the aging process. IDPs and IDRs are typically characterized by low sequence complexity; their sequences are enriched in hydrophilic residues and depleted of hydrophobic residues. [50]. Although such peculiarities of the amino acid sequences preclude IDPs/IDRs from spontaneous folding, the resulting structural plasticity confers multiple functional advantages on these proteins such as multifunctionality and binding promiscuity [8, 10, 14, 20, 51–54]. This dramatically broadens the sequence-function relationships in proteins. Moreover, it represents a departure from the classical “lock-and-key” model of protein enzymatic activity as well as the “induced fit” model of protein-partner interaction, [55] toward the more general “protein structure-function continuum” model. In the continuum model, a given protein exists as a dynamic conformational ensemble that contains multiple proteoforms characterized by a wide variety of structural features and possessing various functional potentials [13, 56–58]. IDPs also usually contain molecular recognition features (MoRFs) — disorder-based protein-partner interaction sites that acquire structure as a result of binding to partners [18, 59–63].

An illustrative example of an important IDP related to aging that has MoRFs and other characteristic IDP features is MYC proto-oncogene protein (a.k.a. c-MYC, UniProt ID P01106). The OSKM Yamanaka cocktail for epigenetic reprogramming is composed of MYC together with Oct-4 (Q01860), SOX2 (P48431), and KLF4 (O43474); each of which contain multiple IDRs (Figure 1a-d). MYC is a transcription factor; it is also a chromatin remodeling protein that regulates cell proliferation [64, 65] and a central player in oncotransformation [66]. Moreover, MYC modulates telomerase catalytic unit (TERT) expression [67] (telomere attrition aging hallmark) and is in use for cells reprogramming [68] and organism rejuvenation [69] (intervention on epigenetic alterations aging hallmark, however, excluded in most recent rejuvenation studies [70, 71]). MYC can also interact with a variety of partners due to its IDRs (Figure 1d). IDRs in general are known to contain MoRFs (marked by yellow segments, Figure 1e), which in turn are known to contain regions capable of binding-induced folding. Indeed, numerous examples of regions capable of binding-induced folding, that can be found in PDB structures, are located within MoRFs, IDRs, or in sequence regions with a PONDR-FIT [72] disorder score around 0.5 (Figure 1e).

**Figure 1.**
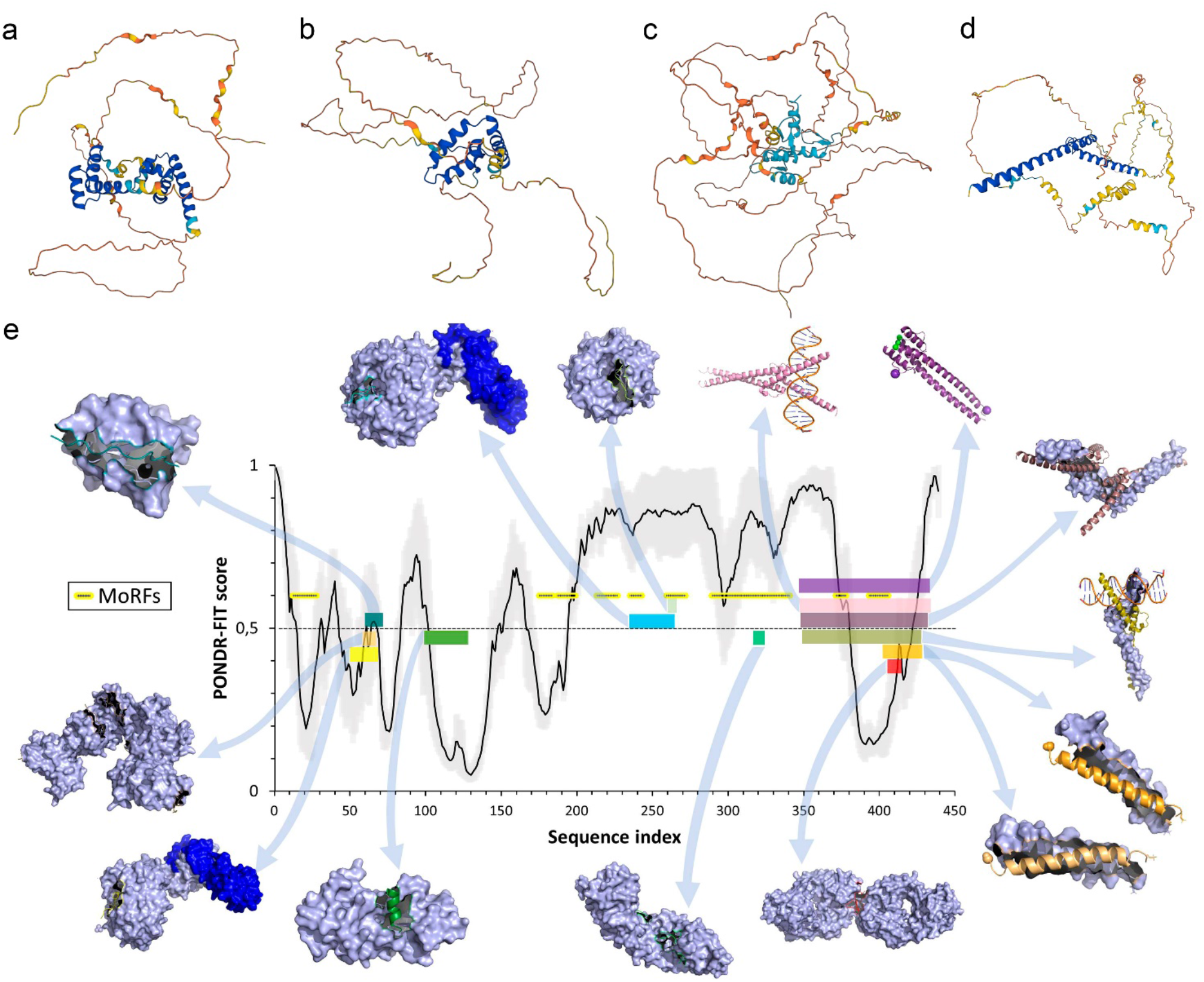
Illustration of structural disorder of OSKM Yamanaka cocktail constituents. A-D. AlphaFold prediction of structures for Oct-4 (Q01860), SOX2 (P48431), KLF4 (O43474), MYC (P01106). Structure prediction reveals presence of IDRs (unstructured regions). AlphaFold structures match partial experimental structures inside ordered regions. E. MYC interaction with different binding partners illustrates peculiarities of one-to-many signaling. An intrinsic disorder prediction by PONDR-FIT algorithm on the MYC amino acid sequence is shown in the center of the figure (above 0.5 threshold = disordered regions, down = ordered, grey – errors) along with the structures of various regions of MYC bound to 14 different partners. The various regions of MYC are color coded to show their structures in the complex and to map the binding segments to the amino acid sequence. Starting with the Fbw7-Skp1-MycNdegron complex (bottom, left, yellow MYC protein, two blue partners, pdb 7T1Z), and moving in a clockwise direction, the Protein Data Bank IDs and partner names are as follows for the 13 complexes: (6C4U – FHA with Myc-pTBD peptide), (1MV0 – BIN1), (7T1Y –Fbw7-Skp1), (4Y7R – WDR5), (5I4Z – apo OmoMYC), (5I50 – double-stranded DNA), (6G6K – MAX bHLHZip), (1NKP – DNA), (1A93 – MAX HETERODIMERIC LEUCINE ZIPPER), (2A93 – MAX HETERODIMERIC LEUCINE ZIPPER), (2OR9 – anti-c-MYC antibody 9E10), (1EE4 – YEAST KARYOPHERIN (IMPORTIN) ALPHA), and (6E24 – TBP-TAF1).

The disorder content of an organism’s proteome increases with the complexity of the organism, which indicates the importance of IDPs [73–75]. Such prevalence of intrinsic disorder is explained by its important functions. First, the conformational transition occurs through intermediate disordered states. Second, a number of proteins require a transition from a structured state to a disordered state in order to perform a function. In addition, some proteins do not fold into a specific 3-D structure, but remain in the form of a conformational ensemble. In this case, some functions will be feasible precisely due to the complete absence of structure.

To the best of our knowledge, there is no systematic analysis linking aging-related proteins to their structural disorder, propensity for liquid-liquid phase separation (LLPS) and aggregation. It is also worth annotating biological processes, signaling pathways, and protein-protein interaction networks according to their enrichment in IDPs. To fill this gap, we present here bioinformatics analysis of 1624 proteins, selected from different ageing-related databases, studies and networks to evaluate the prevalence of intrinsic disorder in these proteins associated with aging. We also show that signaling pathways associated with aging contain IDPs on different hierarchical levels. Enrichment in IDPs and LLPS-prone proteins was shown for the proteins sub-group involved in DNA-associated aging hallmarks. Some aging-related IDPs were classified as “dangerous”, based on the prediction of their propensity to aggregation and, in the same time, to LLPS. Knowledge of the level of intrinsic disorder and the set of functions of the proteins under consideration will make it possible to deepen the understanding of the mechanisms of aging and outline ways to slow it down.

## 3. Methods

### 3.1. Sources: proteins associated with aging

Two different approaches have been implemented to collect a set of aging-associated proteins. The first approach is based on the selection of proteins from databases, such as GenAge (version 20, 307 genes with experimental evidence of direct relation to human aging), Gene Ontology (GO:0007568 term ‘aging’, 277 proteins), and KEGG (Longevity Regulating Pathway, 91 proteins) [76–79].

The second approach is to consider proteins whose expression profile changes with age. It is worth to note, that it was shown that expression of some proteins could increase in middle age relative to young age level and decrease in the end of life (see Supplement Note 1).

#### Proteomics review (Blood): 337 proteins

We used datasets with significant change in the level of protein expression during aging in blood plasma (altered intercellular communication aging hallmark, parabiosis intervention) (collection from [80], proteins that were found in 3 or 4 or more articles – a total of 337 proteins).

#### Gene expression review: 284 proteins

In a study by Marjolen Peters (2015) and colleagues, a large-scale analysis of gene expression in human peripheral blood was carried out. The selection of genes differentially expressed with aging was carried out in two stages. At the first stage, 2,228 genes were selected using literature analysis [81]. At the second stage, 7,909 more blood samples were analyzed, which confirmed the differential expression for 1,479 genes, of which 897 are negatively associated with chronological age, and 600 are positively correlated with it. This analysis was performed using blood samples from people of European descent. To distinguish between changes caused by cellular composition and biological processes, clustering was carried out for groups of genes negatively and positively associated with age using GeneNetwork. For our analysis, only those proteins that remained after clustering were selected – 184 genes that showed a decrease, 100 genes that showed an increase.

#### Chaperome: 163 proteins

Proteostasis maintenance system is of big importance for the aging viewing here through proteome point of view. The study by Mark Brehme and colleagues (2014) showed that the expression of chaperones strongly changes with age: the level of expression of 32% of chaperones corresponding to ATP-dependent molecular machines decreases, and 19.5% of chaperones, corresponding to ATP-independent ones, increases [82]. For this analysis, 332 genes were selected, of which 88 are chaperone genes, 244 are co-chaperones. Experimental analysis of the expression of 48 samples of the superior frontal gyrus taken from healthy individuals, whose age ranged from 20 to 99 years. Analysis of the correlation with aging showed the enrichment of two clusters with certain functional families of chaperones. TPR (tetracopeptide repeats) containing proteins tend to induce expression, while HSP40 does the opposite. In our study, an analysis was carried out for groups showing different directions of expression with aging.

For our analysis, only those proteins with age-related changes were selected – 101 genes that showed a decrease, 62 genes that showed an increase.

#### Ageing brain: 418 proteins

In a study by Lu et al. (2004), age-dependent regulation of gene expression in the human brain was studied [83]. The analysis was carried out on samples of the frontal lobe of the brain of 30 individuals from 26 to 106 years old. Two clusters of identically regulated genes were identified – decreasing and increasing expression with age. Interestingly, the group of people under 42 years of age showed the most homogeneous expression pattern, as did the group over 73 years of age. The expression of genes responsible for synaptic function and the plasticity required for memory and learning was significantly reduced in the aging brain. The same can be said about neuronal signaling, calcium homeostasis, vesicular transport. On the contrary, genes regulating the stress response and DNA repair mechanisms were induced. For this study, 418 genes changing expression with age unambiguously, were selected.

Thus, a set of proteins, one way or another related to aging, was assembled. It claims to reflect the overall picture due to different sources and approaches to data collection.

For our analysis, only those proteins with age-ralated changes were selected – 263 genes that showed a increase, 155 genes that showed an decrease.

#### GWAS of centenarians: 35 proteins

In a study by Yi Zeng and colleagues of 2,178 Han Chinese centenarians, 35 proteins and 4 processes were identified that were significantly associated with lifespan [84].

#### Full aging proteome: 1624 proteins

The complete aging proteome analyzed in this study includes 1624 proteins with experimental evidence. 307 GenAge + 277 GO Aging + 91 KEGG LRP + 337 Proteomics review (Blood) +284 Gene expression review (184 DOWN +100 UP) + 163 Chaperome (62 UP + 101 DOWN) + 418 brain (263 UP and 155 DOWN) + 35 GWAS centenarians = 1912, but actually it was 1624 due to intersections.

In this whole Aging proteome, 446 proteins were DOWN-regulated, 394 proteins were UP-regulated, and 784 were not assigned as UP- or DOWN-regulated. Assignment was done by intersecting the information about gene transcription or proteome analysis by three works, such as Gene expression review, Chaperome, Ageing brain, listed in the 3.1. Sources: proteins associated with aging.

#### Random Proteins Control: 400 proteins

As a control sample, 400 human proteins were randomly taken from the UniProt database [85]. The control sample does not overlap with the protein sets described above.

#### ML aging related

As a comparison, machine learning (ML)-derived dataset of aging proteins was used [86]. Another approach to the analysis of the correlation of intrinsic disorder and proteins involved in aging processes was implemented based on a study by Chab Kerepesi and colleagues (2018). In this study, human proteins were classified as related/not related to aging using machine learning methods. The models were built on the basis of 21,000 feature descriptions of proteins obtained using the UniProt [85], Gene Ontology, GeneFriends databases. The proteins present in GenAge were considered as a selection of aging proteins, and all other Swiss-Prot proteins as non-aging proteins. The models were trained using 5 iterations of cross-validation: all analyzed proteins were divided into 5 non-overlapping samples, 4 of them were used for model training, and 5 for testing. As a result, numerical values ranging from 0 to 1, were obtained for all proteins [86]. These values correspond to the expected relevance of a protein to aging. Based on the obtained model, new proteins that might play key roles in the aging process were identified. The results of the final model are in agreement with the previously known fact that human proteins associated with aging tend to primarily interact with each other.

We considered these proteins independently. We found that only 184 proteins of our dataset are assigned as related and 1440 as unrelated to aging by this approach.

#### Enzymes, Ordered Protein Control: 507 proteins

Enzymes, with their capability to catalyze specific reactions, are typically considered as ordered proteins. 507 enzymes from the collected dataset of 1624 proteins were extracted by UniProt search of proteins with EC (enzyme category) numbers and were used as ordered protein control.

### 3.2. Disorder Prediction: Description and Comparison of Algorithms

The usage of several computational tools to predict the level of protein intrinsic disorder for a given sequence is common practice today. This is because different tools use different algorithms to make predictions and different features, such as amino acid composition, content of charged residues, hydropathy, etc., to describe proteins. As a consequence of these differences, disorder prediction algorithms often show considerable variation in their results. For a comparison of the features of different predictors see Supplementary Note 2. Until an ideal reference algorithm and feature set is established for disorder prediction it will be necessary to consider the phenomenon of protein intrinsic disorder from multiple perspectives. In doing so, a deeper understanding is gained, and a more complete picture is formed.

In this work, we used Rapid Intrinsic Disorder Analysis Online (a.k.a. RIDAO, https://ridao.app) [87], a newly developed platform for disorder analysis that yields results from PONDR VLXT [88], PONDR VSL2 [89], PONDR VL3 [90], PONDR-FIT [72], and IUPred2A [91] in a single run. RIDAO also performs CH-CDF analysis [92] using PONDR VLXT. Each of these predictors is well established and trained on datasets emphasizing various aspects of protein intrinsic disorder. The PONDR predictors employ in this work utilize neural networks while IUPred2A is a physics-based model that does not rely on machine learning.

Each algorithm takes the FASTA formatted amino acid sequences provided to RIDAO as input and outputs a number between 0 and 1 for each amino acid in the input sequence. If the predicted value for a residue exceeds or equals 0.5, this residue is considered disordered. RIDAO averages the obtained per-residue values for the whole protein sequence to provide a mean disorder score (MDS) that describes the level of protein flexibility as a whole. The predicted percent of intrinsically disordered residues (PPIDR) is defined as the number of disordered residues divided by the length of the sequence. Based on their levels of intrinsic disorder, proteins are classified as highly disordered (MDS ≥ 0.5, PPIDR ≥ 30%), moderately disordered (0.25 ≤ MDS < 0.5, 10% ≤ PPIDR < 30%) and ordered (MDS < 0.25, PPIDR < 10%) [93]. The MDS is not directly related to PPIDR (in particular, at 100% PPIDR value of MDS can be anything in the range from 0.5 to 1); therefore, these two methods of protein intrinsic disorder evaluation should be analyzed using correlation.

Important information regarding global classification of proteins based on their levels of intrinsic disorder can be obtained utilizing binary predictors such as charge-hydropath (CH) plots [50, 94] and cumulative distribution functions (CDFs) [94]. By combining these classifiers, one can discriminate proteins in four structurally different classes [92, 95, 96]. The corresponding CH-CDF graph is generated as follows: the y-coordinate is calculated as distance from the boundary in the CH plot, and the x-coordinate is calculated as the average distance of a corresponding CDF curve from the CDF boundary. Positive and negative x-values correspond to ordered and disordered (by CDF) proteins, whereas positive and negative y-values correspond to intrinsically disordered and ordered proteins (by CH), respectively.

On the resulting CH-CDF graph, the lower-right quadrant (Q1) corresponds to ordered and compact proteins (agreement of both predictors); lower-left quadrant (Q2) comprises of proteins predicted to be disordered (by CDF) but having compact conformation (by CH). These proteins usually are native molten globules, or hybrid proteins, containing considerable. Q3, the upper-left quadrant, includes proteins, which are disordered according to both predictors: native coils and native pre-molten globules. The upper-right quadrant (Q4) is usually poorly filled. These rare proteins are predicted to be disordered by CH but ordered by CDF.

Notably, in this work, disorder was not calculated for proteins from the collected datasets that were too short (< 25 residues) or for which a FASTA sequence could not be obtained from UniProt. Additionally, because the disorder prediction algorithms used herein are not designed to operate on non-canonical amino acids, selenocysteine was replaced with cysteine when present. Since for some GeneIDs, there are several human proteins, a reviewed UniProtID entry was used.

### 3.3. Alpha-MoRF prediction

In this study, MoRF regions were predicted using the ANCHOR web server [97], which is available at http://anchor.enzim.hu/index_multi.php. Amino acid residues with a value greater than 0.5 were considered as MoRF regions. MoRF, disordered regions visualized for many proteins in the D^2^P^2^ database [98] (available at https://d2p2.pro/search).

### 3.4. Aggregation propensity prediction

Aggrescan server [99] (http://bioinf.uab.es/aggrescan/) was used for aggregation propensity assessment for the each protein. FASTA formatted sequence was used for calculations.

Two output parameters used for the aggregation propensity prediction – NnHS and Na4vSS. Hot Spots are pieces of protein, which are usually inaccessible to the solvent and are hydrophobic, but when the protein undergoes partial unfolding, they crawl out and provoke aggregation. Aggrescan uses amino-acid aggregation-propensity values (propensities to be Hot Spots) from the experiments [100]. Hot Spots (HS) is defined as a Region with 5 or more residues on sequence with an a4v larger than threshold (HST) and no proline (aggregation breaker)). Used NnHS parameter is a normalized number of Hot Spots for 100 residues (number of Hot Spots divided by the number of residues in the input sequence and multiplied by 100)

Normalized a4vSS for 100 residues (Na4vSS, a4vSS divided by the number of residues in the input amino-acid sequence and multiplied by 100, where a4vSS is a Sum of amino-acid aggregation-propensity value, averaged over a sliding window).

Protein is assessed as aggregation-prone if it has positive or low negative Na4vSS value or has high NnHS (number of Hot Spots) [99].

### 3.5. LLPS prediction

To perform the prediction of probability for liquid-liquid phase separation (LLPS) for the proteins two algorithms were used: PSPredictor [101] (http://www.pkumdl.cn:8000/PSPredictor/) and FuzDrop [102–104] (https://fuzdrop.bio.unipd.it). FASTA formatted amino acid sequence was used as input, proteins were predicted as LLPS positive whet value was ≥0.5 for PSPredictor and ≥0.6 for FuzDrop.

### 3.6. Disorder parameter choice: Correlation matrix

The correlation matrix of disorder predictors assessments (Figure S1) and other properties of all studied proteins represents score (Pearson standard correlation coefficient), equals 1 if two values are fully correlate. Following parameters bring new independent information: ‘CDF’ (Cumulative Distribution Function), ‘PONDR-FIT PPIDR’ (PONDR^®^ Fit assessment of predicted percentage of intrinsically disordered residues), ‘IUPred2A(Short) PPIDR’ (assessment of predicted percentage of intrinsically disordered residues by IUPred2A(Short) predictor), ‘VLXT PPIDR’ (PPIDR estimated by PONDR^®^ VLXT predictor), ‘VL3 PPIDR’ (PPIDR estimated by PONDR^®^ VL3 predictor), ‘VSL2 PPIDR’ (PPIDR estimated by PONDR^®^ VSL2 predictor), ‘CH’ (Charge-Hydropathy value), ‘Percentage of MoRF Residues’ (percentage of residues prone to fold under interaction with the partner), ‘Protein length’ (size of the protein in number of residues).

On the base of this analysis PONDR-FIT, CH, CHF scores were selected as representative disorder evaluations.

### 3.7. Data processing and statistical analysis

1. The prediction and analysis of protein intrinsic disorder was carried out using the disorder analysis platform, RIDAO [87] (https://ridao.app).
2. The PANTHER web server was used to classify the assembled sets of proteins according to biological processes. Automated obtaining of the proteins UniprotIDS, Fold enrichments, p-values, GO terms from xml files was carried out using the Python libraries (see Supplement Notebooks). All 1624 proteins from Aging proteome were analyzed by PANTHER [105]. Overrepresentation Test (Released 20221013) on GO biological processes complete [10.5281/zenodo.6799722] for *Homo sapiens*, reference dataset was whole human proteome. Statistical Fisher’s test with False Discovery Rate (FDR) correction was used with significance level 0.05. Fold enrichments is number of proteins in BP of interest for the Aging Proteome divided by expected number of proteins in BP of interest for the random dataset of the equal size as Aging Proteome. nFoE – normalized fold enrichment.
3. To compare the samples, the Wilcoxon test was used, due to the fact that the distribution of proteins according to the level of internal disorder is not uniform. The Wilcoxon test is nonparametric and tests the null hypothesis that the medians or means of the populations from which two or more samples are taken are identical.
4. Bootstrapping was used to quantify the differences between the medians of the compared samples of significantly different sizes. Bootstrapping is a method of determining statistics by generating pseudo-samples of same size. In our study, bootstrapping was carried out to determine the confidence interval, which is the difference between the median values obtained by generating random subsamples of compared data sets (without returning).

### 3.8. Analyzing IDPs in Interactomes: CytoScape-assisted STRING PPI network creation and assessment of GO Biological Processes, Cellular Compartment enrichment by Bingo app, Panther and STRING Functional Enrichment

Protein-protein interaction (PPI) networks were built in Cytoscape (v.3.9.1; www.cytoscape.org) [106] by following protocol. CytoScape program with STRING App [107] and BINGO [108] applications was used. For PPI-network creation STRING protein query search was done for the whole collected proteins dataset. Then, created network was analyzed (tool “Analyze network”) to calculate number of edges for the nodes (proteins). Defined in such a way, Degree value was used for the determination of hub proteins.

Calculated as previously described (Methods 3.2) proteins disorder scores were added to the Node descriptors table by function Import with key column selected by query name. Excess of edges (protein-protein interactions) was removed by change STRING score to 0.95 and subscores textmining, databases, experiments to 0.25, coexpression to 0.03, cooccurence, neighborhood, fusion to 0. Nodes clustering was done by Scaling (Layout Tools Scale=8) and Edge-Weighted Spring Embedded Layout (clustering by score, i.e. by interaction confidence (distance matrix obtained from the String global scores, so interacting proteins with an higher global score have more chances to end up in the same cluster). Network was drawn with node size reflecting hubness (degree), color – intrinsic disorder state (according to RIDAO calculation of PONDR-FIT values), edge color as betweenness to visualize the most confident interactions in bolder manner.

Finally, Biological Process (BP) or Cellular Compartment overrepresentation (enrichment) analyzes were done by BINGO application for proteins subsets from clusters. For the verification that enriched process is really describe selected cluster STRING Enrichment also was calculated by STRING Functional Enrichment. Double verification was done by clusters analysis in Panther [105] (http://www.pantherdb.org/).

## 4. Results

Bioinformatics analysis of structural intrinsic disorder of aging-associated proteins revealed the functional importance of full and partial absence of the structure.

### 4.1. Design of the analysis

A brief scheme of the study design is visualized in Figure 2. As discussed in the Methods section 3.1., a dataset of 1624 proteins related to aging was collected utilizing two different approaches: searching the Aging Databases and literature overview. For each protein of the resulting dataset, its intrinsic disorder levels were analyzed using predictors of the PONDR family. Next, these proteins were classified by disorder into distinct groups: ordered, partial IDPs and IDPs. Using PANTHER [105] and BinGO [108], key biological processes, overrepresented among the obtained dataset, were identified and further analyzed for IDPs enrichment. KEGG pathways related to aging was analyzed for presence of IDPs at different hierarchical levels. Following this step, protein-protein interaction (PPI) network of all selected aging-related proteins were extensively analyzed using Cytoscape software (for clustering) with BINGO application (for Biological Processes and Cellular Compartment Enrichment) [108] along with PANTHER and STRING Functional Enrichment analysis of protein clusters.

Afterwards, propensity for aggregation (by Aggrescan) for IDPs with high propensity of liquid-liquid phase separation (LLPS, by FuzDrop and PSPredictor) were assessed to find dangerous proteins prone to toxic aggregation.

**Figure 2.**
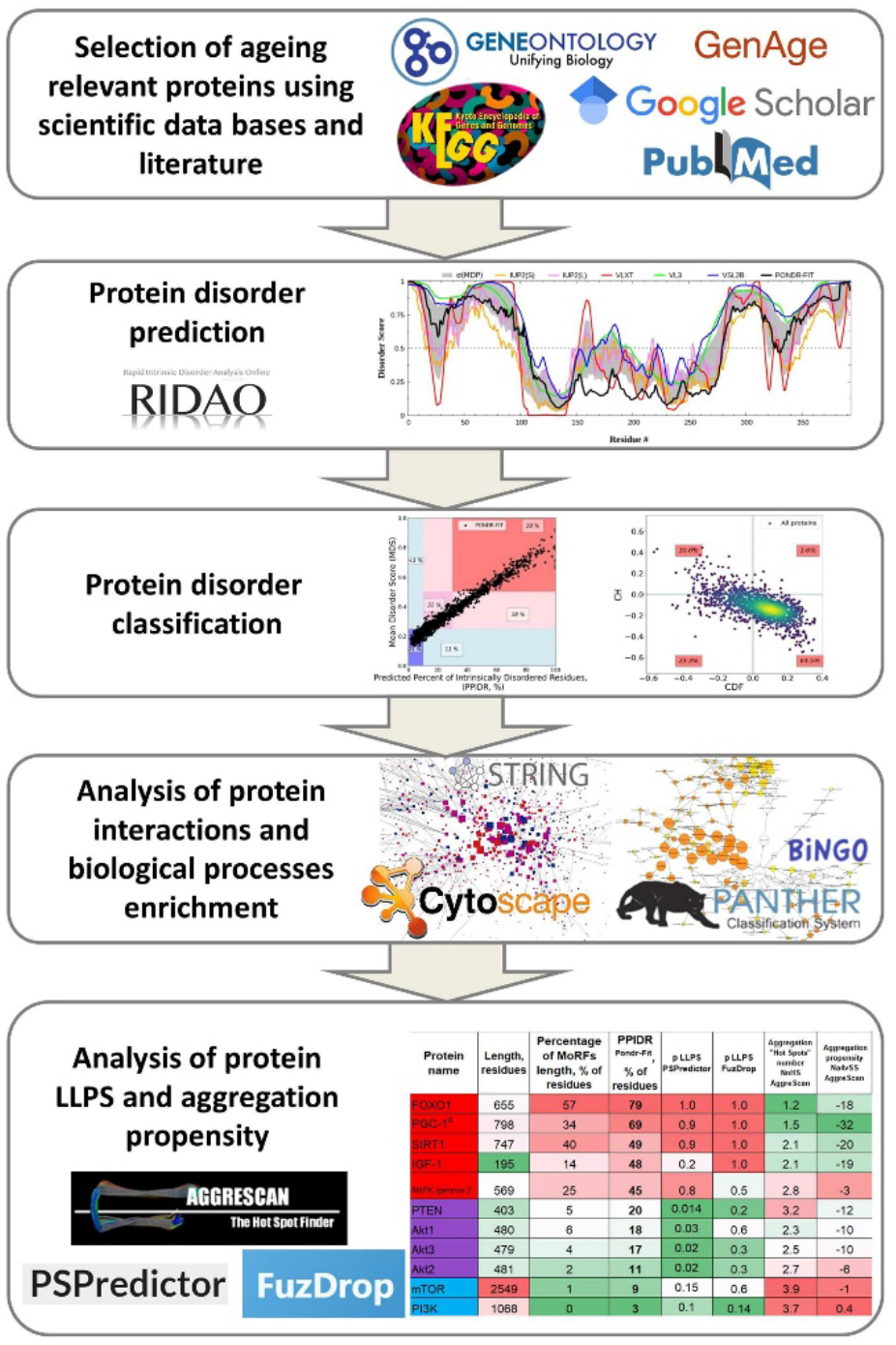
The scheme illustrating the study design.

### 4.2. Disorder evaluation results

Protein structural disorder evaluation by agreement of machine-learning based predictors was done by PONDR-FIT meta-predictor. Combination of counting disordered residues (predicted percent of intrinsically disordered residues (PPIDR) score) or averaging disorder score per-residue values for the whole protein sequence (Mean Disorder Score (MDS)) (Figure 3a) reveals that the majority of the selected proteins are moderately or highly disordered, 36% of proteins in the dataset are highly disordered and could be called IDPs (18% of proteins are in pink area and 18% of proteins are in the red area where PPIDR ≥ 30%). Because PPIDR and MDS PONDR predictions show agreement between each other, and PPIDR representation was used for the intrinsic disorder calculation in further parts of the study (PPIDR^PONDR-FIT^). CH-CDF analysis, based on physical properties and machine-learning prediction, also assigned around 36% of proteins as IDPs (Figure 3b, dots on the left side from vertical threshold CDF=0).

Whole human proteome has slightly higher amount of IDPs by PPIDR^PONDR-FIT^ (Figure 3d, 18% and 21.5% of proteins are in the pink and red areas) and CH-CDF analysis (Figure 3e, around 40%). So, there is no large difference of IDPs amount in our aging proteome in comparison with whole one.

Thus, examination of aging proteins dataset, collected from genomic and proteomic databases and studies, for protein level of intrinsic disorder showed a high percentage (about 35%) of highly disordered proteins that is on the same level as for whole human proteome (around 40%) (Figure 3c, f, according to threshold PPIDR^PONDR-FIT^ ≥ 30%).

**Figure 3.**
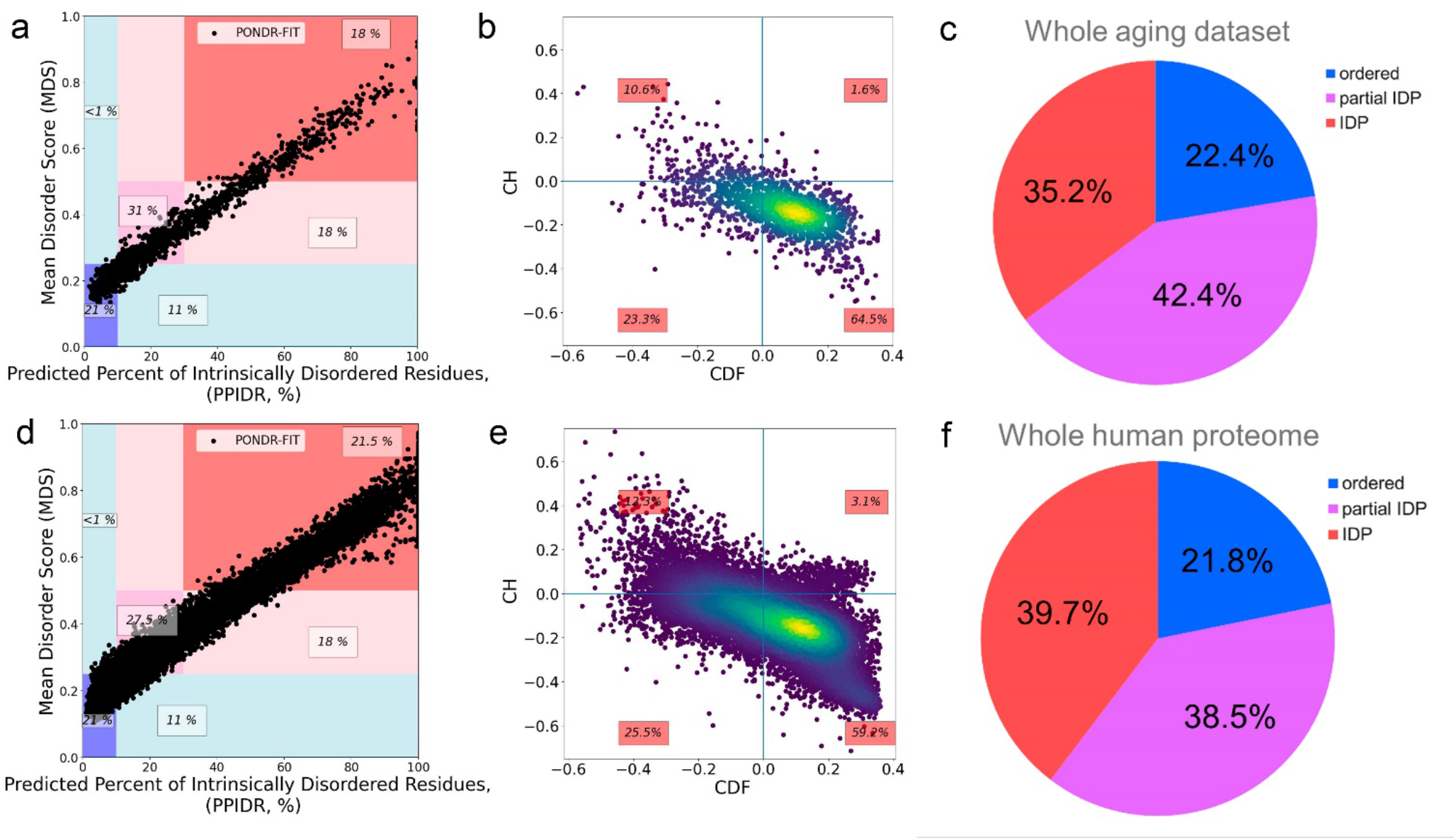
Analysis of disorder scores for proteins from whole dataset. A, D. MDS^PONDR-FIT^ (PPIDR^PONDR-FIT^) plot for Aging proteome and whole human proteome respectively. Labeled regions are marked by 0.25 and 0.5 values of MDS and 10 and 30% of PPIDR. MDS (Mean Disorder Score, describes the level of protein flexibility as a whole) and PPIDR (predicted percent of intrinsically disordered residues) scores from PONDR-FIT assessment correlates with each other. B, E. CH-CDF analysis for Aging proteome and whole human proteome. CH-CDF analysis shows IDPs as agreement of CH and CDH prediction (upper-left quadrant) and as only CDF-predicted as disordered (bottom left quadrant). C, F. IDPs abundance of Aging proteome and whole human proteome.

### 4.3. Analysis of the aging-associated proteins for LLPS propensities

Since intrinsically disordered proteins are prone to toxic aggregation (β-amyloid, α-synuclein, tau protein, and others [108–117]), it is necessary to analyze if highly disordered aging-associated proteins are potentially dangerous as drivers of proteinopathies. To achieve this goal, it is important to evaluate, as first step, the probability of liquid-liquid transition for proteins (Figure 4). Analysis of the LLPS propensity was done using algorithms (FuzDrop, PSPredictor) that has not high correlation (Figure S2a-c) between each other and with disorder score, so could be considered as independent.

**Figure 4.**
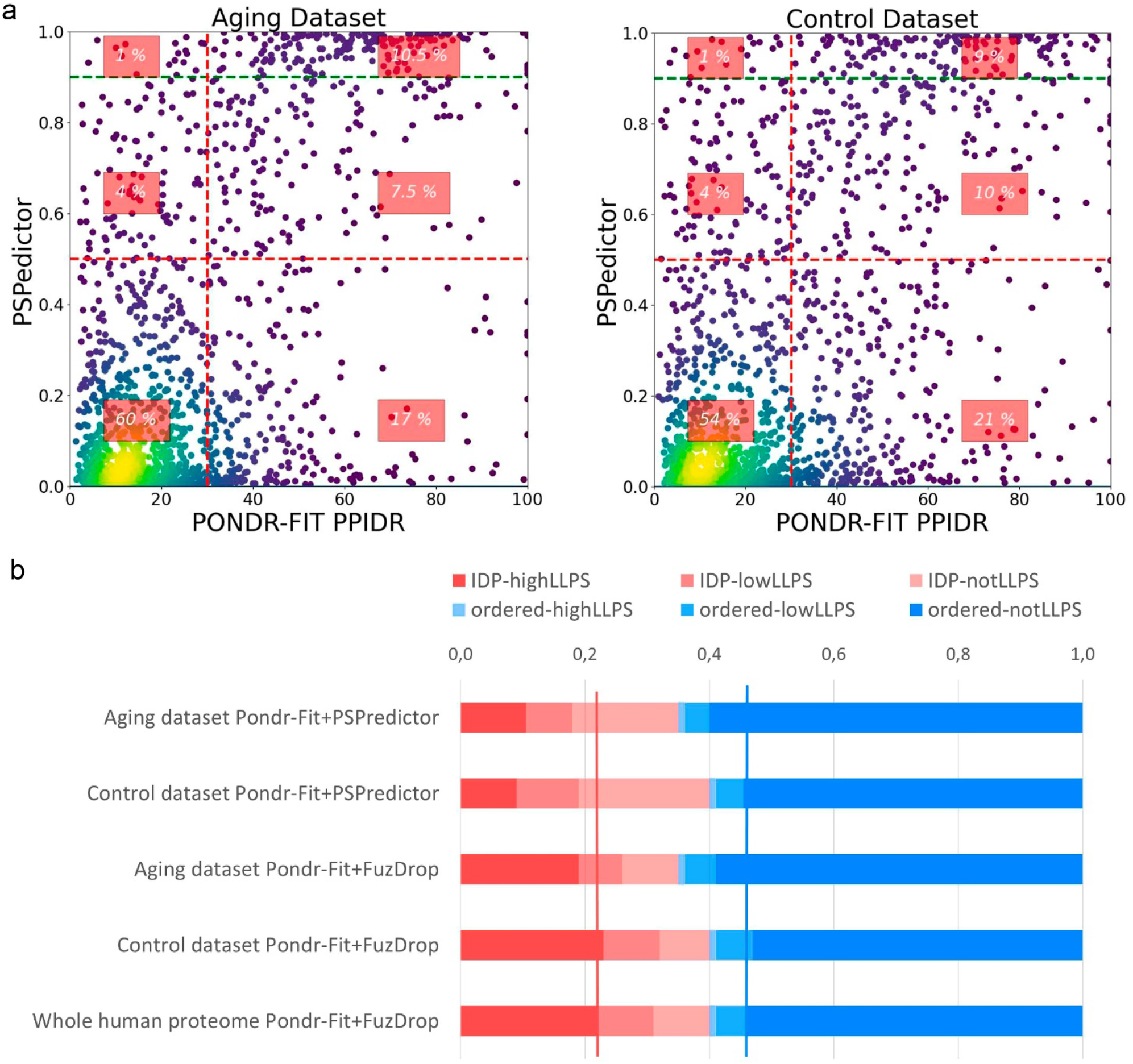
Correlation between intrinsic disorder and LLPS propensity for aging dataset VS controls. A. PSPredictor score in relation to PPIDR^PONDR-FIT^ for aging dataset (left panel) VS control dataset of random 1624 proteins (right panel). B. Comparison of partitions of proteins on groups by disorder and LLPS propensity. Red columns are for IDPs, densest red is for LLPS-prone IDPs, less dense red is not LLPS-prone IDPs. Blue columns are for ordered proteins, densest blue is for not LLPS-prone ordered proteins, less dense blue is for LLPS-prone ordered proteins. Control dataset (that was used with PSPredictor) shows results which is quite similar to whole human proteome (from FuzDrop).

It is of interest to analyze distribution of proteins in space spanned by PPIDR^PONDR-FIT^ and LLPS propensity predictors, PSPredictor and FuzDrop. Using PSPredictor, it is possible to compare the collected ageing dataset and control set of random 1624 proteins (Figure 4a). 60% of proteins from the aging dataset (against 54% for the control set) are placed in the lower-left region corresponding to ordered proteins with low probability for LLPS. Similar observation can be done using larger, whole proteome dataset with FuzDrop predictor (Figure 4b, blue bars for Aging dataset are higher than for controls and whole proteome, see also Figure S2d for FuzDrop LLPS(PPIDR^PONDR-FIT^) plots).

We can conclude from Figures 3 and 4 that one of a specific characteristics of the collected ageing dataset is elevated percentage of ordered proteins with low probability for LLPS. There is no over-abundance of intrinsic disorder in aging dataset. However, it is well known, that enzymes and secreted proteins, that are simple to monitor and overrepresented in all studies, are biased to ordered state. Furthermore, if we could find special IDP-enriched biological process, it would be quite useful for anti-aging interventions. To accomplish that, we turned to more detailed analysis.

### 4.4. Biological Processes: Analysis of the level of disorder in various biological processes

We checked median disorder level for the protein subsets from different sources, which constitute our full aging dataset, independently (Figure 5a, see detailed sources description in Methods): databases (KEGG Longevity Regulating Pathway, GeneAge, Gene Ontology term Aging), for proteins, which expression profile changes with age, selected by means of literature analysis (Proteomics Review, Gene Expression Review, Chaperome, Aging Brain), a control sample (random dataset of 400 human proteins), aging-associated and non-aging-associated proteins according to the Csab Kerepesi classification of proteins [86] (ML Aging-related, ML Aging not-related), and ordered proteins control (Enzymes).

Comparison of distributions of intrinsic disorder in each sample with control random dataset of 400 proteins, representing median proteome level of disorder, have shown that neither of sources have median disorder higher than control. As a positive control, highly ordered Enzymes dataset showed strong statistical difference (P ≤ 0.0001) according to the Wilcoxon test between sample and control. Reviews summarized significantly different expressed proteins with aging (Proteomics and Gene Expression Reviews). Proteomics and Gene Expression Reviews are united in investigation of proteins (translated or transcribed) in blood plasma. Thus, the reason behind lower medians is possibly due to the fact that the majority of proteins in these two datasets are secreted ones. Three available aging-related datasets (in Figure 5a, ‘GenAge’, ‘Gene Ontology Aging’ and ‘Longevity Regulating Pathway’), have a distribution very similar to that of the control sample.

Figure 5a also compares proteins that are classified based on the machine learning model [86] as related and not related to aging, that are not taken into account during building of our aging proteome, collected on the basis of experimental evidence. The Wilcoxon test confirms the differences between ML-related and ML-not related to aging samples. This result to some extent supports the hypothesis that for aging-related proteins, there is a tendency to a higher level of intrinsic disorder. However, this model usually classifies proteins with a high number of interacting partners as ‘aging-related’, feature that is inherent for the IDPs [118]. Thus, this result might be due to the specifics of the model and might not represent the general picture. Furthermore, level of median disorder for ML aging-related sub-dataset is as high as in Control sample. So, no disorder abundance for the ML aging-related sample. Further, we did not use ML-derived aging dataset.

To deeper investigate relations between intrinsic disorder and aging, we selected proteins (from the our aging dataset) for which the direction of expression change was known, using information from literature [81, 82, 119]. UP- and DOWN-regulated proteins did not show statistical differences in levels of intrinsic disorder from the control sample and each other (Figure S3). However, both UP- and DOWN-regulated proteins sets are comprised of proteins, which take part in a number of important biological processes. These biological processes could differ in their average levels of intrinsic disorder. Therefore, it is more important to analyze which of the most pronounced processes (in the resulting dataset) have increased (or decreased) levels of disorder, and if aging proteins are more disordered among all proteins of a single considered process.

**Figure 5.**
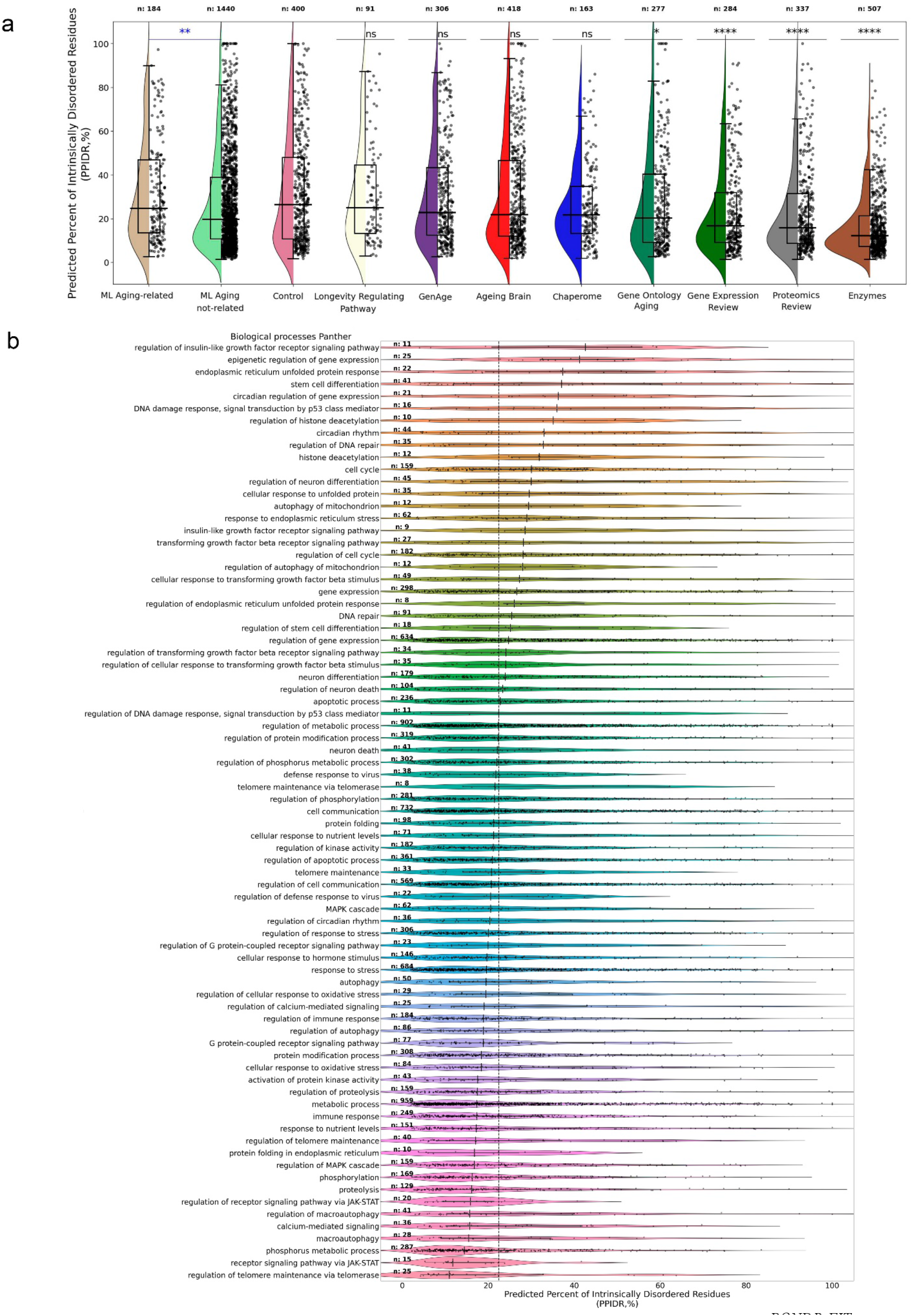
Distribution of aging-related proteins to BP and disorder score (PPIDR^PONDR-FIT^). B. Aging-related processes, enriched for our aging dataset, with distribution of disorder score for the proteins.

An analysis of the total set of proteins on biological processes was carried out using the PANTHER web server [105], which showed the enrichment of many biological processes (BP). Figure 5b shows the distribution of aging-associated proteins among 77 selected BPs, representing aging hallmarks. Processes were selected based on three criteria: relevance to key aging mechanisms [1, 2], enrichment for the BP and regulation of this BP, and the presence of at least 8 proteins from the collected dataset.

All BP sub-groups of aging-related proteins were analyzed to found their disorder distributions (Figure 5b). It could be concluded, that a Gene Ontology analysis of the Aging proteome revealed an abundance of IDPs in one-third of aging-associated processes, especially in regulation of gene expression, proteostasis maintenance, regulation of nutrient sensing, DNA damage response. Such enrichment makes sense due to the known regulatory function of IDPs, need of disorder for interaction with DNA [30], and the existence of hormones in IIS pathway (regulation of insulin-like growth signaling). Another one-third of aging-associated processes are depleted in IDPs, due to the enrichment in enzymes. Finally, one-third of aging-associated processes have IDPs percentage similar to the whole human proteome. Taking into account that for the whole proteome median value of PPIDR^PONDR-FIT^ is 22.44%, it can be stated that among aging-related proteins, mostly ordered proteins take part in the processes below threshold. On the contrary, aging-related proteins of processes above threshold are mostly highly or moderately disordered. Graph also indicates that for the selected aging-associated overrepresented processes intrinsic disorder levels are higher than 10%, meaning that median level of disorder for these processes is higher than the threshold for ordered proteins.

It is of interest to study CH-CDF layout of enriched biological processes. Most of them have similar distribution, with around 40-75% of proteins being in the lower-right quadrant (ordered proteins) (Figure 6b). However, there are BPs with enrichment of ordered proteins by CH-CDF analysis (Figure 6a): receptor signaling pathway via JAK/STAT (100% ordered proteins), regulation of receptor signaling pathway via JAK/STAT (100%), regulation of endoplasmatic reticulum unfolded proteins response (87.5%), phosphorus metabolic process (77.7%), G protein-coupled receptor signaling patway (75.3%), macroautophagy (75%). Contrary, depletion of ordered proteins is shown for other BPs (Figure 6c): epigenetic regulation of gene expression (24% ordered proteins), endoplasmatic reticulum unfolded protein response (31.8%), histone deacetylation (33,3%), regulation of insulin-like growth factor receptor signaling (36,4%), DNA damage response, signal transduction by p53 class (37,5%), circadian regulation of gene expression (38,1%).

**Figure 6.**
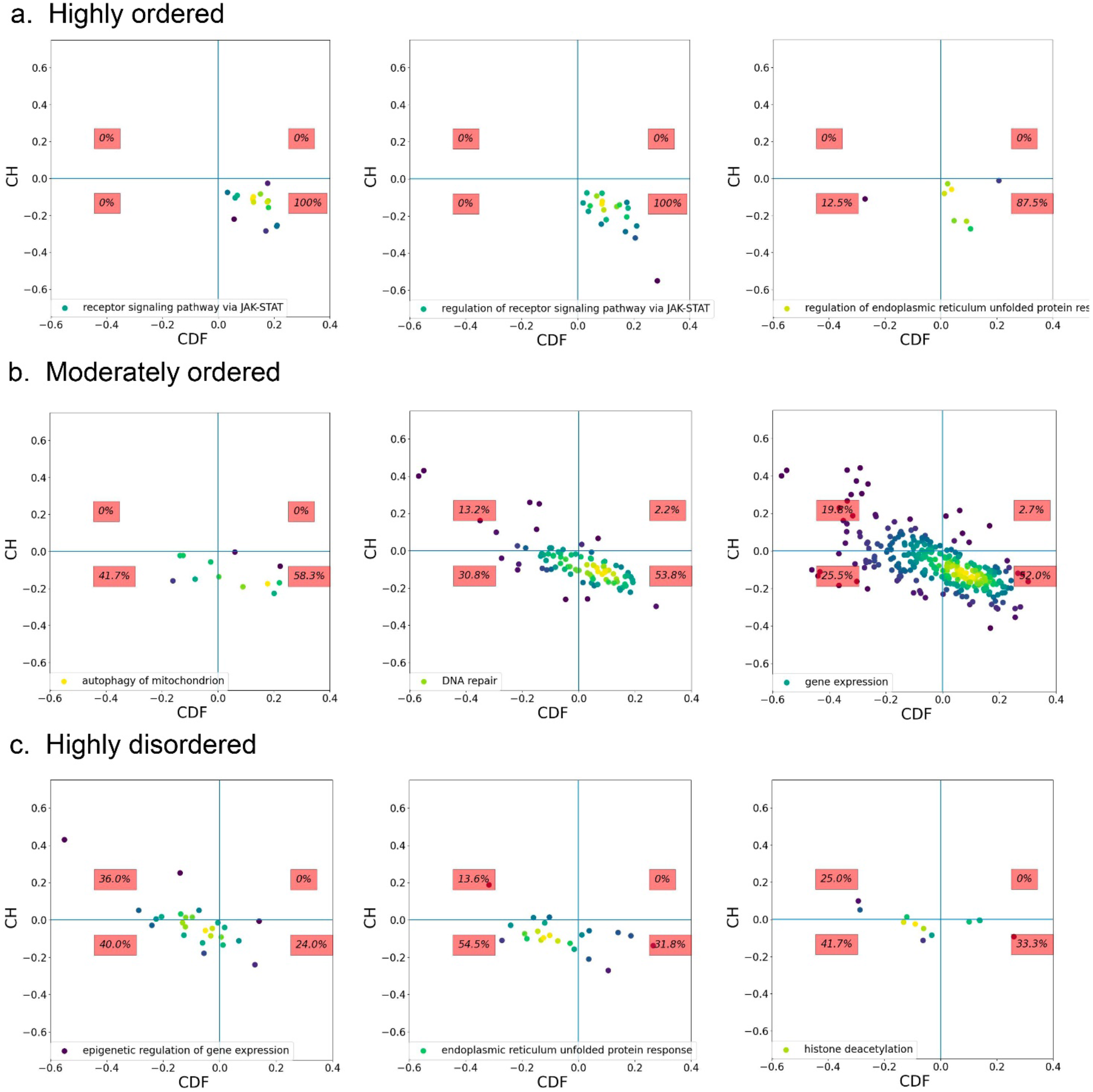
CH-CDF analyses of proteins of our aging proteome in biological processes sub-groups. A. Highly ordered processes (more than 75% of proteins are ordered). B. Moderately ordered processes (number of ordered proteins are between 40 and 75%). C. Maximally disordered processes (less than 40% of proteins are ordered). Three examples for each category presented.

As a result, we could stress, that different approaches of protein structural disorder assessment give consistent results in ranging BPs by disorder (comparison of PONDR-FIT results Figure 5B and same manner plot for CH-CDF Figure S4).

A special distribution in terms of intrinsic disorder may or may not be a characteristic of the proteins from the collected data set in comparison to other proteins for each BP. To test this hypothesis, three aging-related processes were selected, the proteomes of which included a large number of proteins from the obtained data set for the correctness of the statistical analysis. For each process, the distributions of intrinsic disorder from the obtained dataset and the remaining proteins of this particular biological process were compared (Figure S5). Results show that intrinsic disorder can be not only higher for ageing-related proteins, but also lower. Therefore, the presence and absence of protein intrinsic disorder may be important for aging-related protein subgroup within the biological process.

Once the overrepresented in the resulting dataset BPs were identified, it is needed to see the impact to the enrichment in each BP of three disorder groups: ordered, partial IDP, and IDP. Two complementary estimations by Panther (Figure 7 and Figure S6, sorted by IDP impact into enrichment) and BinGO (Figure S7, consistent with Panther results) conducted. Bаsed on these analyses, highly disordered proteins are most widely represented in the following processes: regulation of gene expression, histone deacetylation, endoplasmatic reticulum unfolded protein response, and regulation of insulin-like growth factor receptor signaling pathway. Ordered proteins are depleted in DNA repair, cellular response to oxidative stress, and others, in which IDPs enriched in comparison with whole proteome.

**Figure 7.**
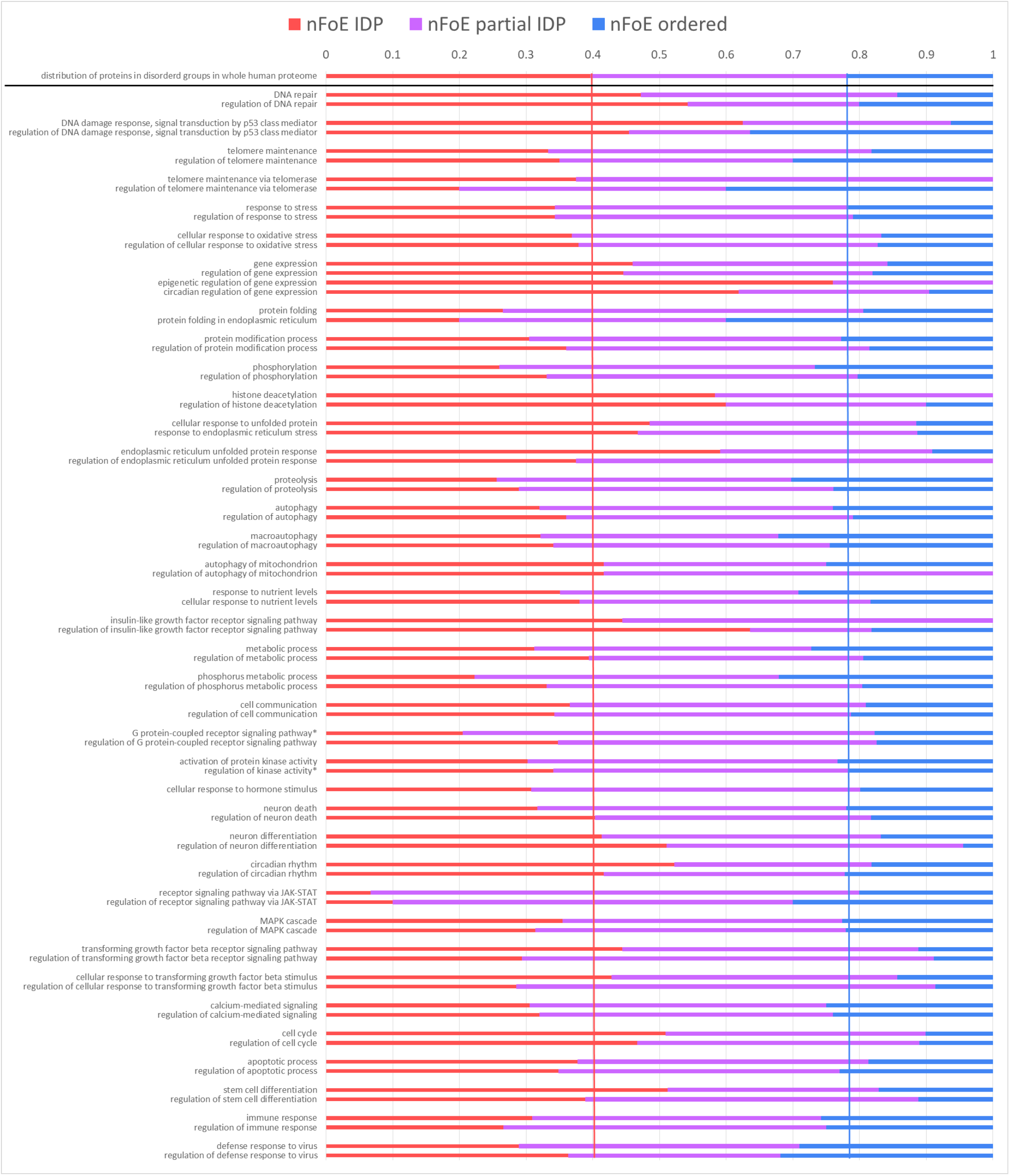
Impact in BP enrichment of proteins with different disorder assignment. PANTHER Overrepresentation Test (Released 20221013) collects enriched biological processes. Ones, associated with aging hallmarks, were chosen. All 1624 proteins from Aging proteome were analyzed on GO biological processes complete [10.5281/zenodo.6799722] for Homo sapiens organism, reference dataset was whole human proteome. Statistical Fisher’s test with False Discovery Rate (FDR) correction was used with significance level 0.05. Impact of different disordered classes to enrichment represented by coloring: blue – ordered, violet – partially disordered, red – IDP. For the comparison of IDP presence in each BP with expected level for the whole proteome, there is shown distribution of whole human proteome proteins on IDP-partial IDP-ordered groups and vertical thresholds for IDP and ordered proteins presence drawn. Plots presents normalized Enrichment, not-normalised FoE plots shown in Figure S7. nFoE – normalized fold enrichment (number of proteins in BP of interest for the Aging Proteome divided by expected number of proteins in BP of interest for the random dataset of the equal size as Aging Proteome). * values for the “regulation of kinase activity” are average of ones for the two GO terms (positive regulation of kinase activity, negative regulation of kinase activity), values for the “G protein-coupled receptor signaling pathway” are average of ones for the three GO terms (phospholipase C-activating G protein-coupled receptor signaling pathway, adenylate cyclase-activating G protein-coupled receptor signaling pathway, adenylate cyclase-modulating G protein-coupled receptor signaling pathway).

Such plots show again that gene expression regulation relies on highly disordered proteins, whereas enzymes-rich processes (such as ‘proteolysis, ‘metabolic process’, etc) are biased toward order, if one would select aging-related proteins from these processes. However, this selection is sometimes ambiguous because there is no universal aging dataset and aging-related BPs. Databases that we used in the present study include different protein sets. Therefore, some incorrect conclusions about biases for order or disorder in aging-related selection can be ascribed to imperfect databases.

### 4.5. The presence of intrinsic disorder in signaling pathways associated with aging

Presence of intrinsic disorder in specific signaling pathways is of big interest due to pathways integrates BPs needed for accomplishing some function. On Figure 8a one can see a diagram of the interaction of key proteins involved in the recognition of nutrients. Distortions in this pathway are one of the hallmarks of aging [1, 2], and interventions against this hallmark being very common [120]. Scheme illustrates growth hormone and insulin-like growth factors signaling (IIS) and dietary restriction (DR) pathways, and the relationship of these pathways and aging. IIS pathway accelerates aging, dietary restriction slows it down.

**Figure 8.**
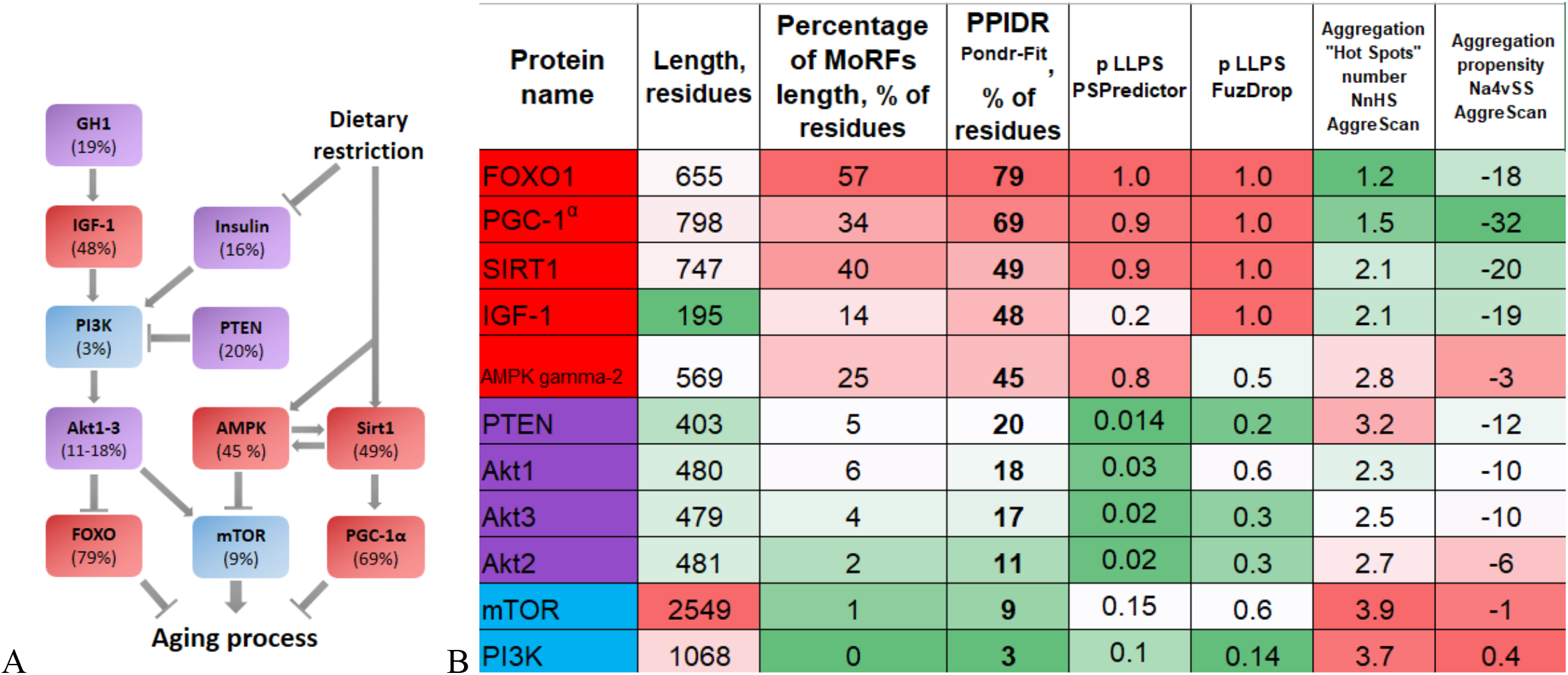
Nutrient sensing signaling pathway of mTOR complex 1. A. Diagram showing key proteins and signaling pathways involved in nutrient recognition. The scheme of the pathway is modified from the publication of Lopez-Otin and colleagues [1]. PPIDR^PONDR-FIT^ values for each protein are labeled in protein boxes, coloring reflects classification by structural disorder: blue – ordered, violet – partially disordered, red – IDP. B. Analysis of the level of intrinsic disorder of key proteins in aging regulating metabolism. Red-to-Green coloring (heat-map) represents high values as Red: long length, long MoRFs length, high disorder score, high LLPS and aggregation propensities. UniProt IDs for the proteins: FOXO1 – Q12778; PGC-1α – Q9UBK2; SIRT1 – Q96EB6; IGF-1 – P05019; AMPK gamma-2 – Q9UGJ0; PTEN – P60484; Akt1 – P31749; Akt3 – Q9Y243; Akt2 – P31751; mTOR – P42345; PI3K – P42336.

The scheme on Figure 8a demonstrates that opposite pathways (IIS, aging acceleration, and DR, aging deceleration) contain IDPs. Figure 8b illustrates the properties of key aging-related proteins in this metabolism regulating network: PPIDR^PONDR-FIT^ as disorder score, abundance of MoRF regions, as well as LLPS and aggregation propensity. We could see that, first, intrinsic disorder is represented differently in key proteins in aging. Second, MoRFs abundance and LLPS propensity correlate strongly with intrinsic disorder, whereas aggregation propensity predominantly anti-correlates with other parameters. Only for MAPK subunit gamma-2 disorder, LLPS and aggregation are high in the same time. Thus, disordered and ordered proteins are mixed in the nutrient sensing signaling pathway.

Another, KEGG Longevity Regulating, pathway is probably the most important pathway for the processes of aging. Analysis of this pathway (Figure 9a) shows that most of the proteins involved in it are moderately (violet coloring) or completely intrinsically disordered (red boxes). The illustration also shows other signaling pathways intersecting with this pathway as ovals. All intersecting pathways are moderately intrinsically disordered – the median PPIDR value fluctuates around 19%. At the same time, for proteins of KEGG Longevity Regulating pathway the median value lies much higher, 25%. Therefore, within this narrow dataset, we can say that aging-associated proteins turned out to be, on average, less ordered.

**Figure 9.**
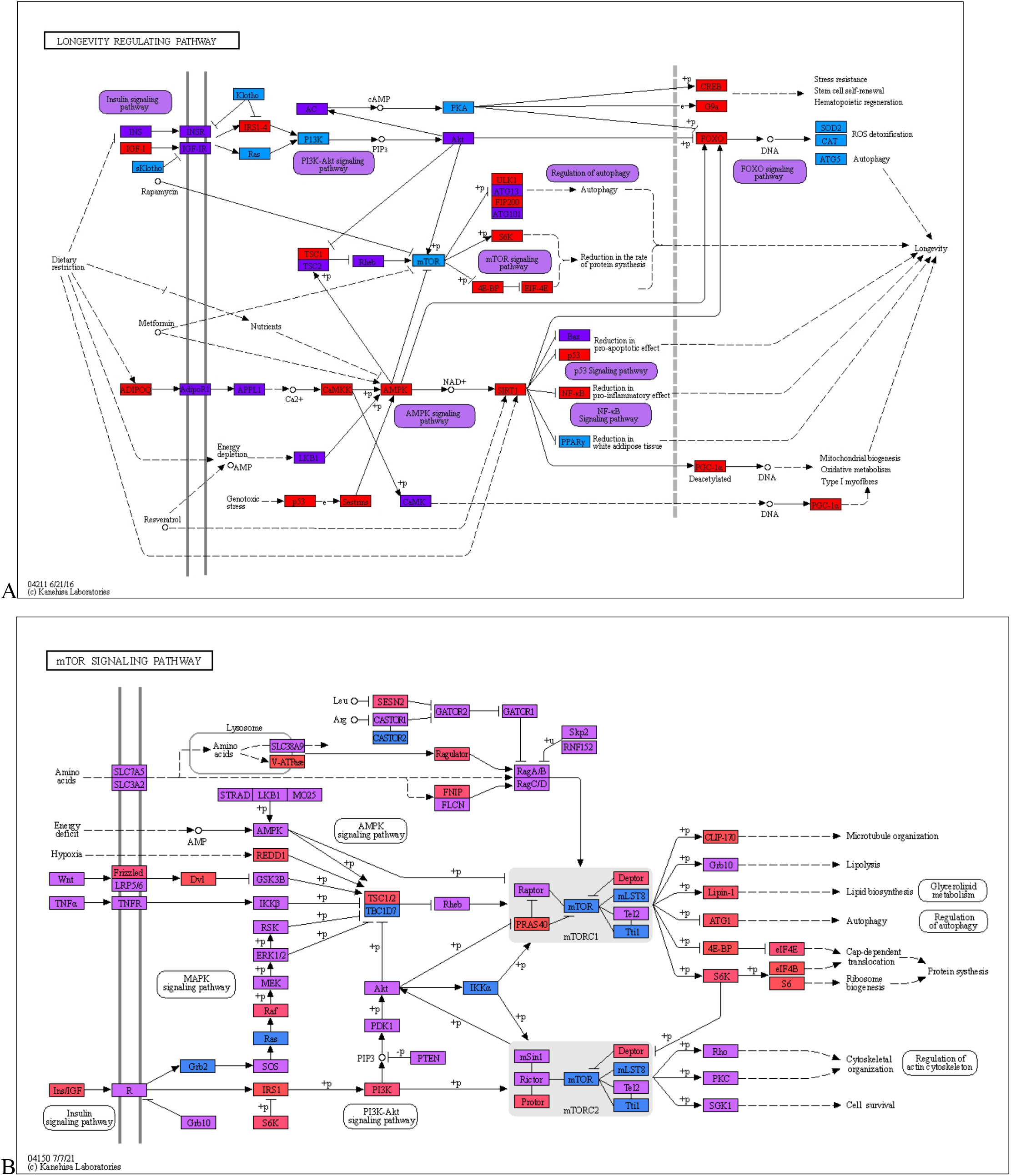
Classification of proteins of the KEGG Longevity Regulating (A) and mTOR signaling pathway (B) according to the level of intrinsic disorder.

It is interesting that both KEGG longevity regulating (Figure 9a) and nutrient recognition (Figure 9b) pathways (which are partly intercepted) have several highly ordered nodes that are followed by 1-2 disordered ones. That looks like brickwork (structure-function continuum [13, 56–58]), where the interaction of ordered enzymes (“bricks”) is controlled by mobile fraction of IDR-containing proteins (“viscous cement”).

Therefore, signaling pathways associated with aging contain IDPs on different hierarchical levels.

### 4.6. Analysis of protein-protein interaction (PPI) networks in BPs

In a study by Yi Zeng and colleagues, 35 proteins and 4 processes were identified as significantly associated with lifespan [84]. The PPI diagram (Figure 10) shows that these proteins and processes are strongly interconnected. On the scheme, the levels of intrinsic disorder of proteins are displayed and colored in accordance with the boundary values – ordered proteins are shown in blue, partial IDPs in purple, and highly disordered proteins (IDPs) in red. From this scheme, it can be concluded that proteins of the MAPK pathway, differently expressed in centenarians, are characterized by the prevalence of intrinsic disorder, since all 4 proteins are highly or partially disordered. The process of immune response can be called moderately disordered, response to stress process is biased to ordered proteins. Thus, 35 proteins inherent for centenarians have different levels of disorder, and proteins of different disorder make different BPs.

**Figure 10.**
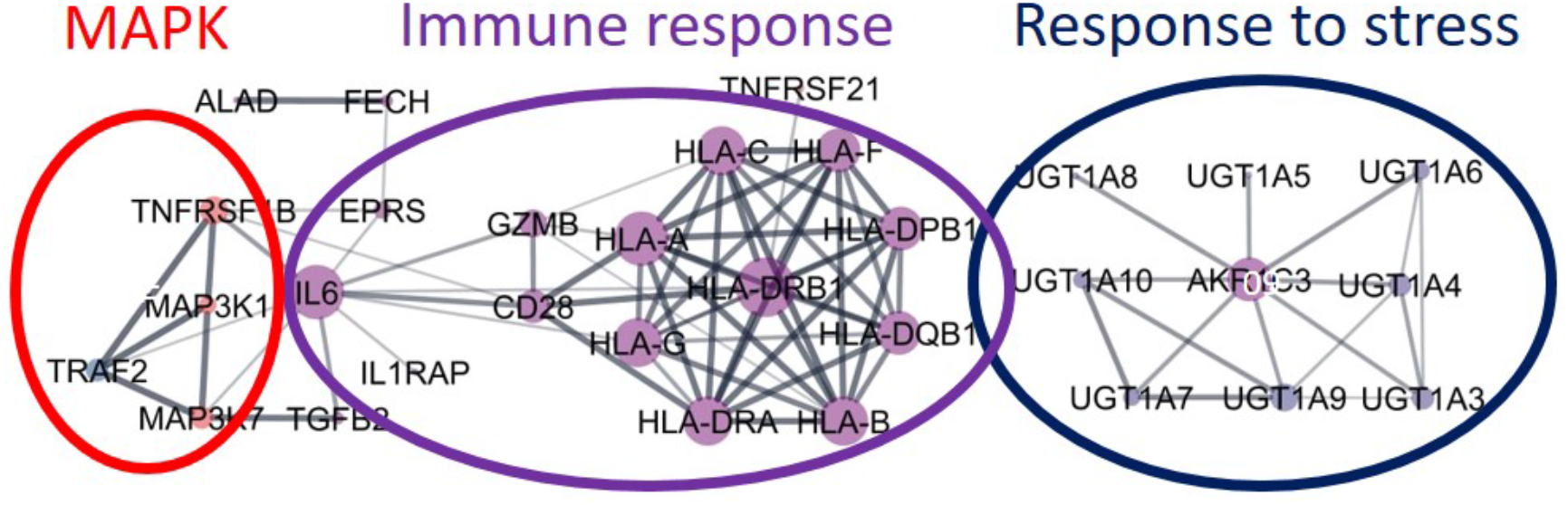
Networks of proteins, characteristic for the Chinese centenarians, colored based on disorder (PPIDRPONDR-FIT).

Similar to this small protein dataset, a study of the various protein-protein networks utilizing STRING database was carried out for our whole Aging proteome. PPI analysis is complementary to the classification by BPs described in previous sections because PPI gives proteins clusters and reveals main functions of studied proteome. Obtained PPI network for the whole aging proteome is not scale-free (degree distribution follows a power-law), i.e. contains clusters. Random protein dataset from human proteome of same size 1624 proteins has no clusters (Figure S8a).

PPI-network of aging-associated proteins was clustered on sub-groups by interaction density. Clusters sharing cellular compartment and biological process, UP- or DOWN-regulation with aging (Figure 11A, assignment of the clusters to cellular compartment and BPs was performed by agreement of Bingo App (Figure S8), Panther (Figure S9) and STRING Functional Enrichment (Figure S10)).

The densest cluster №1, *i.e.* representative one for whole dataset, includes 264 proteins, ∼30% of which are IDPs, nuclear location enrichment, and performs several hallmarks-associated functions: nutrient sensing, intercellular communication, response to stress, maintenance of proteostasis, mitophagy, cellular senescence. In contrast, a much smaller cluster №3 of 131 proteins, enriched with IDPs (∼52% compared to 40% for the whole human proteome, Figure 11B) performs about the same number of functions related to the aging hallmarks, especially DNA-related ones, such as DNA repair, telomere maintenance, epigenetic regulation, stem cells exhaustion, cellular senescence. Such multi-functionality of this cluster is natural due to its enrichment in IDPs. This protein sub-group №3 has high enrichment in “nucleus” location and DNA-associated functions that confirms importance of IDPs for gene expression regulation process, showed in previous sections.

The cluster №4 enriched in the UP-regulated with aging proteins and depleted in IDPs (30% compared to 40% for the whole dataset) regulates immune and stress responses. DOWN-regulated proteins are enriched in cluster №2 ensuring protein folding function. It is also not enriched in IDPs (30%) (see IDPs enrichment on Figure 11B).

Thus, in collected dataset we have UP-regulation of stress responses and DOWN-regulation of structural processes, like proteostasis, which is in consistence with previous works [83, 121].

Another important question is the relation of disorder and characteristics of formed networks. Two different networks were constructed for comparison: one formed by the aging-related ordered proteins (PPIDR^PONDR-FIT^ 0-10%) and another formed by the aging-related highly disordered proteins (PPIDR^PONDR-FIT^ >50%). Network formed by ordered proteins turned out to be nearly 20% less dense and 10% less heterogeneous, than the network formed by the disordered proteins (Figure S11). These results are in line with the idea that intrinsically disordered proteins have more interactions with partners.

LLPS prediction (Figure 11C, Figure S12) showed that IDPs have high probability to be engaged in liquid-liquid phase-transition. Protein clusters associated with molecular hallmarks of aging have LLPS-positive proteins. IDP-enriched cluster №3 has also LLPS-enrichment, so nuclear IDPs associated with aging have high LLPS propensity. These observations indicate that LLPS is coupled to IDP and is quite important for aging.

**Figure 11.**
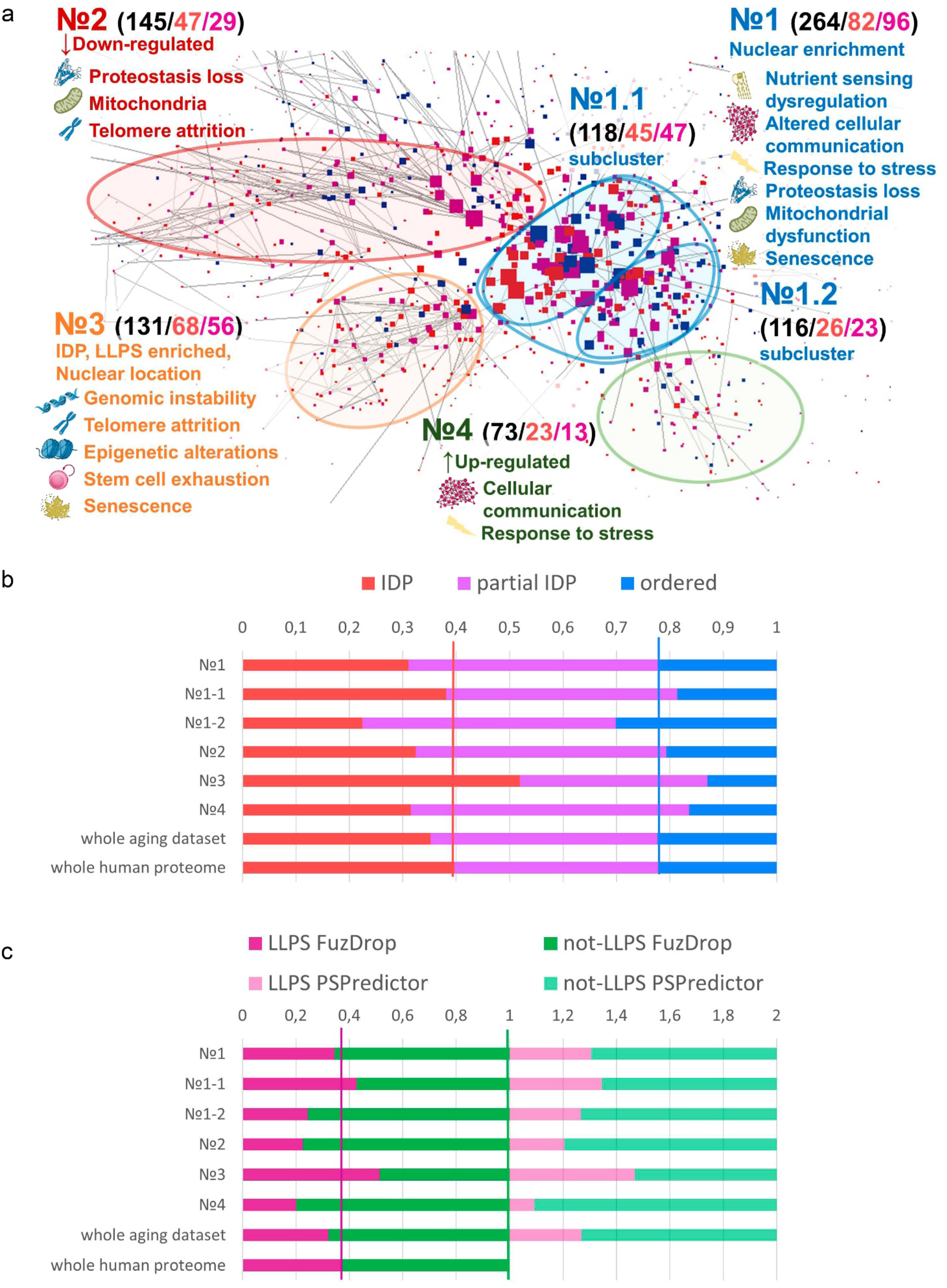
Analysis of PPI network for the full collected dataset of aging-related proteins. A. PPI network showing IDP as red (PONDR-FIT PPIDR > 30%), partial IDP are pink/violet (30%> PONDR-FIT PPIDR > 10%), ordered proteins as blue (PONDR-FIT PPIDR < 10%). Node size is proportional to the number of connections. Four clusters are clearly distinguishable. GO-analysis of clusters proteins subsets (Figure S10, S11) reveals their main functions (BPs) which are attributed to hallmarks of aging and written on the figure. For each cluster total/IDPs/LLPS positive numbers of proteins are shown in black, red, dark pink colors correspondingly. B. Distribution of disorder by clusters, red parts – ordered proteins, violet parts – partially IDP, red parts – IDP C. FuzDrop and PSPredictor results in LLPS-propensity assesment for proteins from PPI network clusters. Violet parts – LLPS – prone proteins, green ones – not LLPS – prone proteins.

### 4.7. In depth analysis of highly disordered proteins from various biological processes

Insoluble protein aggregates are the hallmarks of many neurodegenerative diseases. However, whether aggregates cause cellular toxicity is still not clear, even in simpler cellular systems. Aggregation of TDP-43 is not harmful but protects cells, most likely by taking the protein away from a toxic liquid-like phase [44]. On the other hand, it is recognized that the protein aggregation could be dangerous, *e.g.* aggregated proteins can distort biogenesis and mechanical properties of the proteinaceous membrane-less organelles (PMLO) [47, 122, 123] . Therefore, aggregation and LLPS propensities should be assessed for the IDPs in aging dataset with representative biological functions (BP) and taken into account.

For each of the biological processes found to be significantly overrepresented, some highly disordered proteins were selected for in-depth analysis (Figure 12). It is characteristic for proteinopathies-associated proteins (prion protein, α-synuclein, SOD1, β-amyloid precursor protein, Huntingtin, marked as bold black on Figure 12) that structural disorder, LLPS and aggregation scores are high. For ordinary, not disease-associated, not dangerous proteins, there is usually another picture: ordered proteins have low disorder and LLPS scores but aggregation is predicted to be high, and, vice versa, IDPs have high disorder and LLPS scores but low aggregation. The reasons for such clustering are clear from the structural point of view. Hydrophobic regions inside protein globule of ordered proteins will tend to aggregate with any other protein in the case of misfolding. In other words, structurally ordered domains are often prone to aggregation but they can avoid it if folds quickly [124]. Contrary, IDPs are predominantly hydrophilic and charged, so mostly are not tending to hide any regions from water inside the aggregates. Generally, IDR appearance coupled with both high LLPS and aggregation propensity is characteristic for disease-associated proteins (reviewed in [122]). Therefore, we considered such a combination as a potential danger marker of a protein. For the well-known LLPS positive proteins G3BP1, SERF1, TDP-43 (TADBP), VHL aggregations assessments are close to control proteinopathies-associated proteins. So, we have positive control, checked that simultaneous disorder, LLPS and aggregation reflects dangerous features.

**Figure 12.**
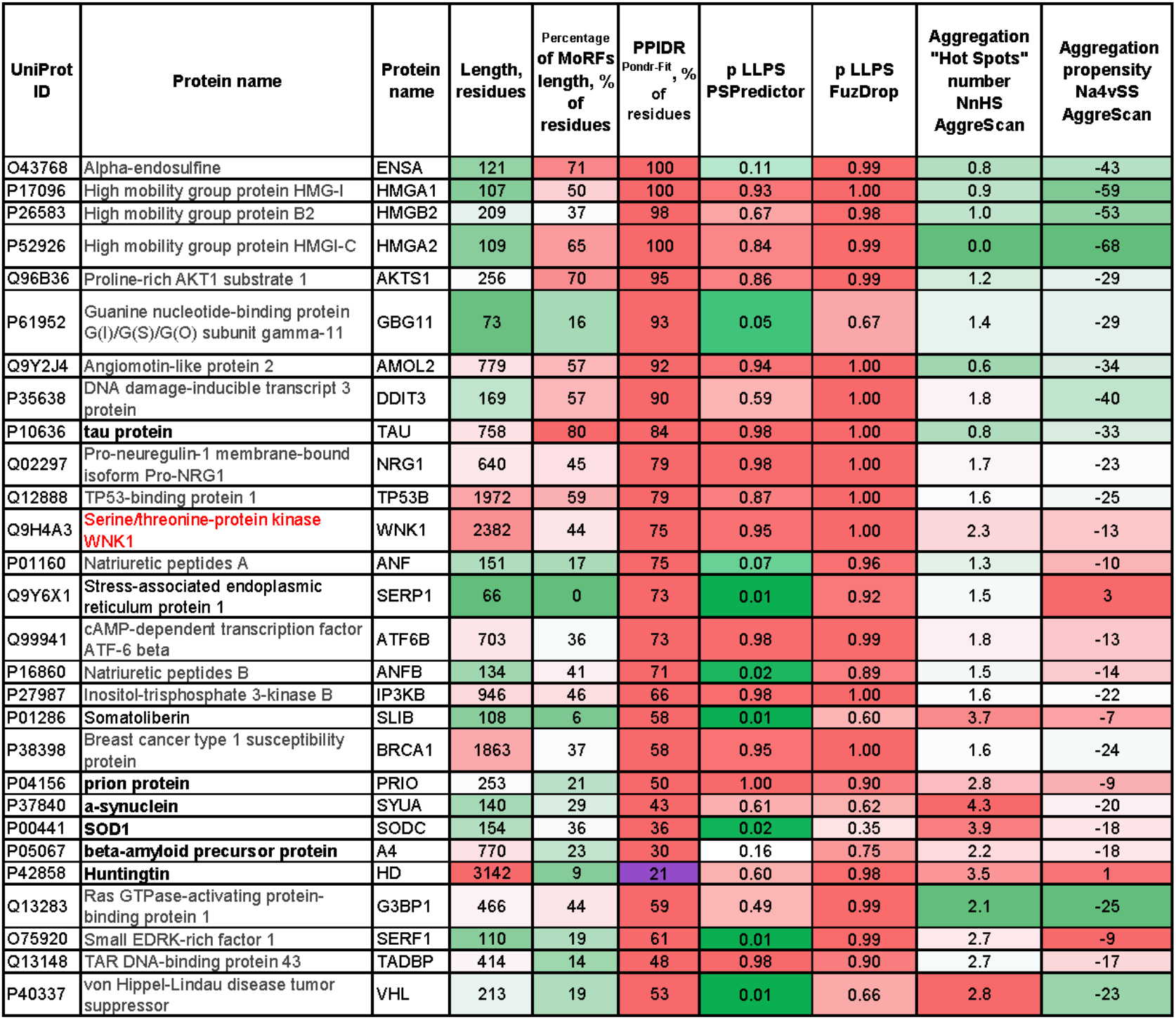
Analysis of the most disordered proteins from different biological processes on “dangerous” features: LLPS propensities and aggregation. Proteinopathies associated proteins highlighted as bold black, new “dangerous” IDP marked as red. Heatmapped columns gradually colored from red to green from top to lowest disorder (MoRFs, PPIDR^PONDR-FIT^), LLPS propencity (PSPredictor and FuzDrop), aggregation scores (AggreScan “Hot Spots” number and area of them).

In the small subset of our ageing dataset, it can be seen that one of the selected proteins, Serine/threonine-protein kinase WNK1 (Q9H4A3), shares “dangerous” characteristics with control diseases-associated proteins. This kinase does form aggregates [125], but has a role of signaling platform, where WNK acts together with other kinases [126]. It is also reported that WNT1 tend to interact with hydrophobic domains of partner proteins “defending” them from abnormal aggregation [127]. WNK1 activity is known to correlate with tumor malignancy [128], but there is currently no evidence that pathologic WNK1 aggregation is associated with the disease. Furthermore, simultaneous high LLPS and aggregation scores were shown at Figure 8b for AMPK gamma-2 (P05496), which is also shown to be capable to aggregation [129].

Based on these observations, we suggest that these two aging-associated proteins could be dangerous in terms of their propensity for toxic aggregation and could cause disruption of proteostasis, but clarification of this issue requires further experimental proof. Therefore, some IDPs, related to aging, were highlighted as “dangerous”, by the prediction of their propensity to LLPS and aggregation.

## Conclusions

IDPs were given little importance before, and they could even be considered as some kind of rudiments that do not have functions as was for non-coding RNA. Recently, the role of both of them has become clearer [130]. Among other things, their significant regulatory role was discovered, as well as the ability to influence the interactions of biopolymers with each other, including aggregation and a more subtle process – liquid-liquid phase separation. LLPS, as it turned out in the last decade, is closely related to the processes of regulation of proteostasis and stress responses. At the same time, the peculiarities of IDPs as well as the presence of phase separation in the cell were ignored for a long time, in particular, in aging research. In order to understand the role of disordered proteins and LLPS in the aging process, we conduct comprehensive bioinformatics meta-analysis. Thus, we worked in the two “dark” places in aging research, both functionally (IDP and LLPS) and methodologically (bioinformatics), the combination of which can provide new information about the aging process.

To better uncover the mechanisms of aging, it is important to understand the prevalence of intrinsic disorder in proteins associated with aging and elucidate which cellular aging-related processes are most enriched in IDPs. Previously, similar intrinsic disorder-centric analyzes have been performed for proteins involved in Alzheimer’s disease [24, 131], Parkinson’s disease [132, 133], as well as frontotemporal lobar degeneration and amyotrophic lateral sclerosis [25, 134]. These works showed presence of IDPs in aging-related diseases.

Based on bioinformatics analysis, we investigated the characteristics of 1624 aging-associated proteins obtained from public databases or published studies. It has been shown that in aging-associated proteins, the disorder and LLPS propensity, coupled to each other, are on average lower than in the control sample and for the whole human proteome. Thus, both ordered and disordered, LLPS positive and negative proteins are important for the aging processes.

Further, aging-associated proteins were attributed to distinct biological processes and consequently to different aging hallmarks. When the Aging proteome was sorted by biological processes, it was discovered that there is significant heterogeneity in the average level of disorder between BPs – about one-third is enriched with ordered proteins and one-third with disordered ones. IDPs are enriched in DNA-associated processes (especially in regulation of gene expression, DNA damage response).

The analysis of signaling pathways showed that IDPs are located at all levels of the signaling system hierarchy – several highly ordered nodes are followed by 1-2 disordered ones. This looks like brickwork (structure-function continuum [13, 56–58], where the interaction of ordered enzymes (“bricks”) is controlled by the mobile fraction of IDR-containing proteins (“viscous cement”). If we add to this that IDP are known for highly specific but weak interactions, we will get even more universality, which allows forming signaling networks of almost any shape, number of nodes and high plasticity for removing or adding new components. It should be further noted that some chains of this pathways are protein complexes consisting of several subunits (such as mTORC1). So, combination of ordered and disordered parts can be observed even within one component of the signaling system, in which one subunit may be responsible for enzymatic activity or scaffold (ordered parts), while other may recruit interaction partners (parts enriched with disorder). That proposes an answer on a very delicate question of how one protein complex (mTORC1, AMPK, Sirt1 and etc.) can quickly interact with dozens or hundreds of proteins, which even may not have a pronounced “lock-and-key” correspondence. The fact that represented core metabolic pathways are highly nutrient-dependent highlighted the degree of human dependence on the environment, which effects on the IDR-containing proteins and LLPS are just beginning to be seriously explored.

For better understanding, aging-associated proteins were grouped by their mutual interactions, and four clusters were discovered. Two of them are mainly localized in the nucleus. One nuclear cluster is enriched with disordered proteins with a high propensity for LLPS and is responsible for DNA-associated functions. Therefore, we confirmed that IDP and LLPS have a special role in aging but for the highest degree within the framework of processes associated with the genome regulation.

UP-regulated proteins with aging, associated with stress response, and DOWN-regulated proteins, performing proteostasis functions, have no difference in intrinsic disorder and LLPS enrichment from each other and from the whole human proteome. Information about expression change with aging for the cluster of disordered and LLPS-prone proteins, enriched in genome regulation, is lacking and of high interest to be explored. Finally, in small subgroup of aging-related proteins that are highly disordered and LLPS positive, we identified two potentially dangerous that are prone to pathologic aggregation – WNK1 and AMPK gamma-2. Literature analysis confirmed their ability to aggregate. We suggest that these two aging-associated proteins may be dangerous in terms of toxic aggregation and proteostasis disruption, but clarification of this issue and analysis for dangerous aggregation in the full collected here aging proteome requires further study.

In summary, our analysis reveals the special role of ID and LLPS in the genetic regulation of aging, highlighting the group of proteins for a more thorough experimental study in that context.

Knowledge of the level of intrinsic disorder and the set of functions of the proteins under consideration will make it possible to deepen the understanding of the mechanisms of aging and outline ways to slow it down. To build robust link between aging and structural disorder, as it was done for other aging hallmarks [1, 2], one should do positive and negative controls – rescue of accelerated aging by expression/depletion of IDPs and premature aging after IDP knock out/overexpression correspondingly. Some studies present so far [135, 136]. Further research is needed to add IDP, LLPS, PMLO disturbance as aging hallmark.

## Abbreviations

IDP: intrinsically disordered protein
IDR: intrinsically disordered region
PONDR: Predictor of Natural Disordered Regions
PONDR-FIT, VLXT, VL3, VSL2, IUPred2A: predictors of protein disorder (existence of the disordered residues in the protein) from PONDR-family
MDS: mean disorder score – average of PONDR scores of each residue for whole protein
PPIDR: predicted percentage of intrinsically disordered residues (with propensity to be disordered higher than 0.5)
CDF: Cumulative Distribution Function
CH: Charge-Hydropathy
MoRF: Molecular recognition features (protein-partner interaction sites that acquire structure upon binding to partners)
LLPS: liquid-liquid phase separation FuzDrop, PSPredictor predictors for LLPS propensity
PPI: protein-protein interaction
BP: Biological Process
Aggrescan: predictor for propensity to aggregation
IIS: insulin and insulin-like signaling
DR: dietary restriction
TERT: telomerase catalytic unit
nFoE: normalized fold enrichment, ML – machine learning

## Statements and Declarations

### Ethics approval and consent to participate

Not applicable

### Consent to publish

Not applicable

### Data Availability

The authors confirm that the data supporting the findings of this study are available within the article and also present as electronic supplementary materials. Additional data will be available on reasonable request.

### Author Contributions

V.D.M and N.S.I. contributed equally to this work. N.S.I., V.I., and V.N.U. conceived and designed study. V.D.M., N.S.I., S.V.N., B.M.G.A., G.W.D, E.V.Z., S.S.M., A.V.F., I.M.K., K.K.T., and V.N.U. conducted research, analyzed data, and collected and analyzed literature data. V.D.M., N.S.I., S.V.N., B.M.G.A., G.W.D, E.V.Z., and S.S.M. designed illustrations. V.D.M., N.S.I., S.V.N., B.M.G.A., G.W.D, E.V.Z., S.S.M., A.V.F., I.M.K., K.K.T., V.I., and V.N.U. wrote the manuscript. All authors have read and agreed to the published version of the manuscript.

