## Supplementary Material for "Chaotic aging: Intrinsically disordered proteins in aging-related processes" (from another version of this preprint: version 1, 2023-04-25)

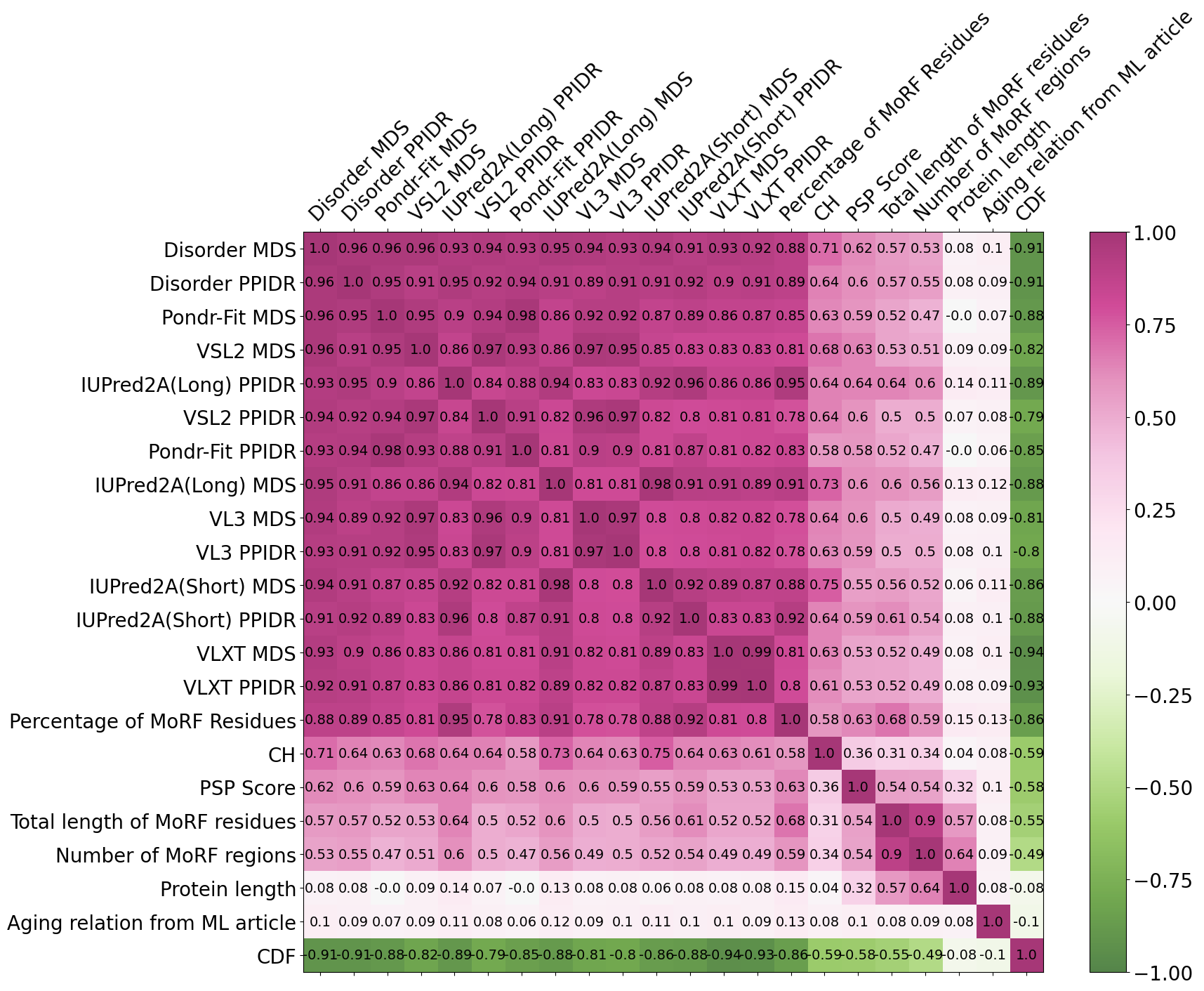

Figure S1. Correlation matrix of disorder parameters for all studied proteins revealing the most representative parameters. Pearson correlation R was used, which is correct because data has multivariable normal distribution.

PONDR - Predictor of Natural Disordered Regions giving propensity to be disordered for each residue, MDS – mean disorder score – average of PONDR scores of each residue for whole protein, PPIDR - predicted percentage of intrinsically disordered residues (with propensity to be disordered higher than 0.5), Disorder – here, result of averaging predictions for 6 algorithms (PONDR–Fit, VLXT, VL3, VSL2, IUPred2A short and long), PONDR–Fit, VLXT, VL3, VSL2, IUPred2A - predictors of protein disorder (disordered residues in the protein), Percentage of MoRF Residues (percentage of residues prone to fold under interaction with the partner), CDF - Cumulative Distribution Function, CH - Charge-Hydropathy, PSP score – prediction to do LLPS (liquid-liquid phase separation) by PSPredictor, Protein length (size of the protein in number of residues), Ageing relation from ML article [86].

Correlation of disorder and LLPS propensity
MF – MoRF percentage, PF – PPIDR^PONDR-FIT^, PSP – PSPredictor, FD – FuzDrop.

**a MF PF PSP FD	MF PF PSP FD**	**MF PF PSP FD**

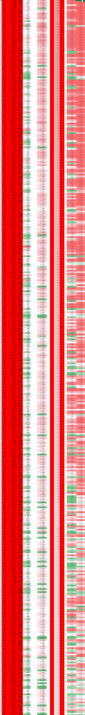

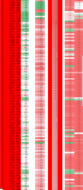

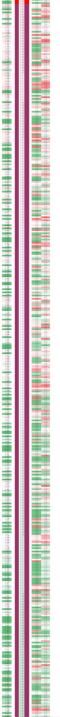

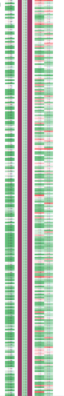

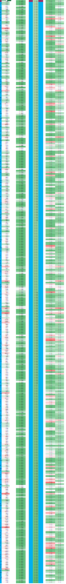

**b**
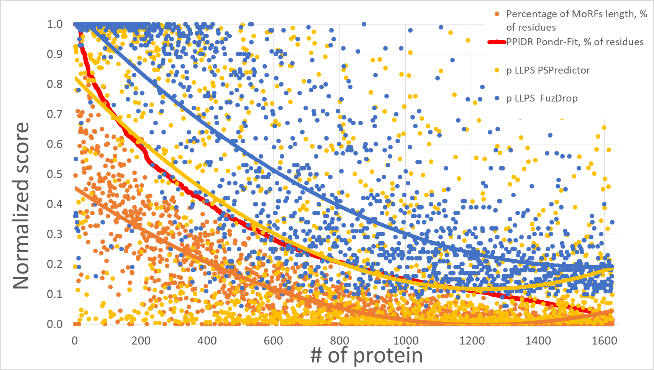

**c**
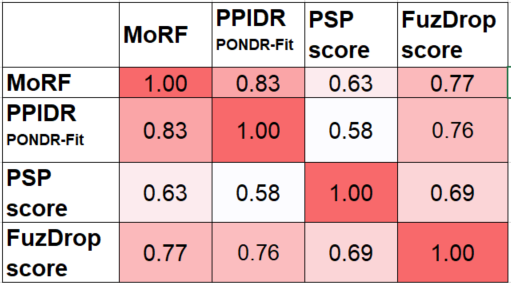

| **d** | Whole proteome | Aging dataset |
| --- | --- | --- |
|  | 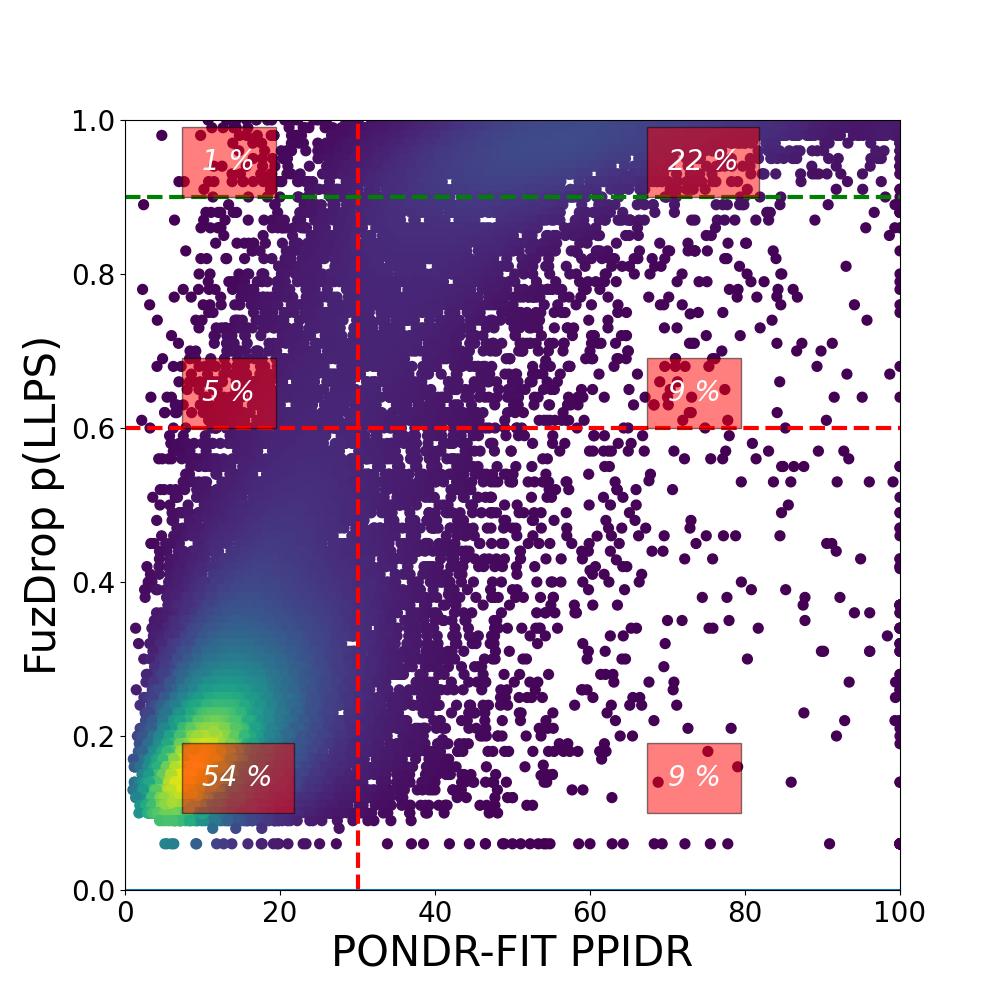 | 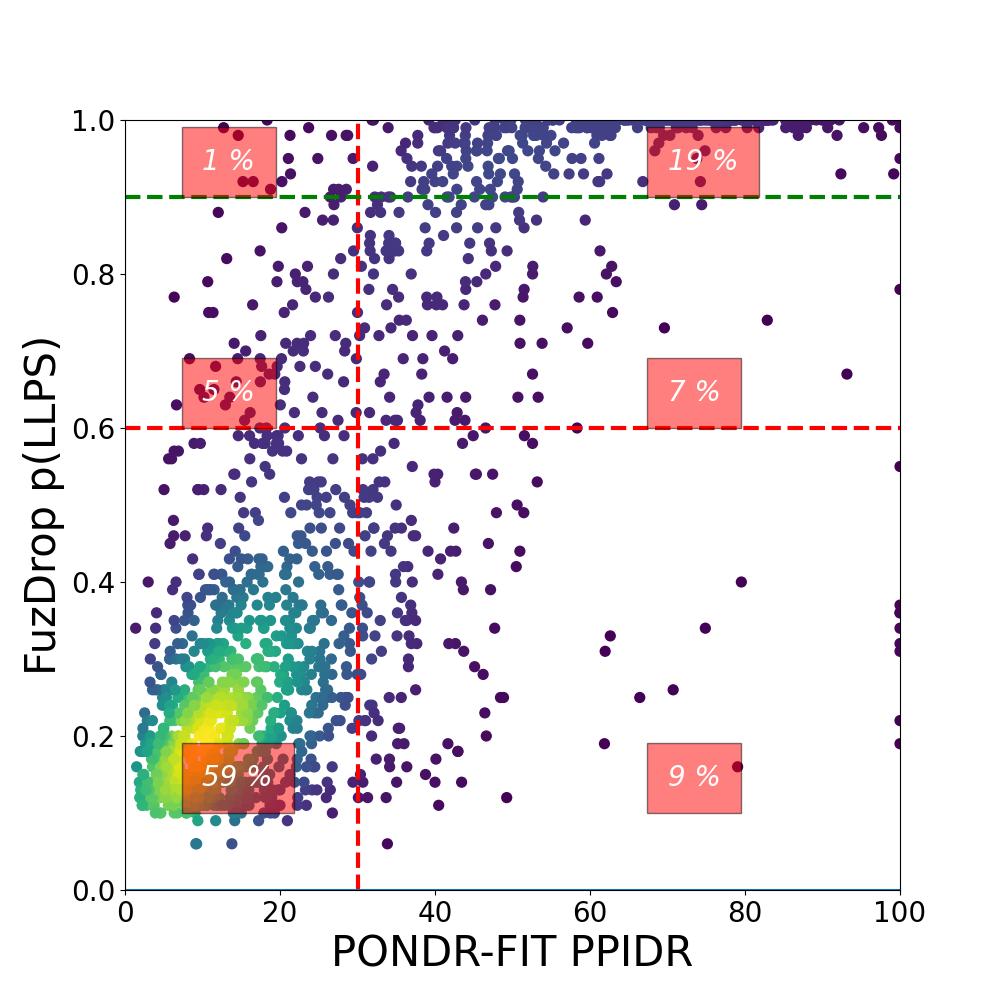 |
|  | Whole dataset | Control dataset |
|  | 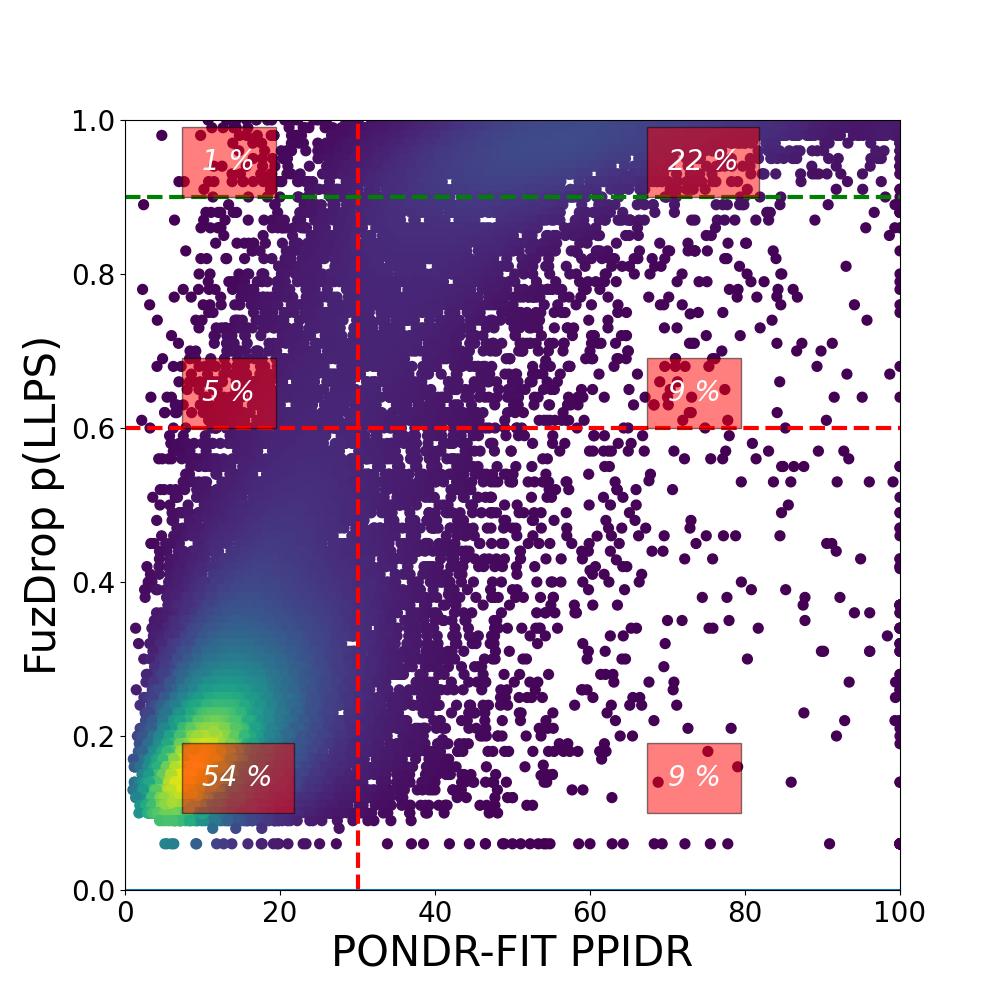 | 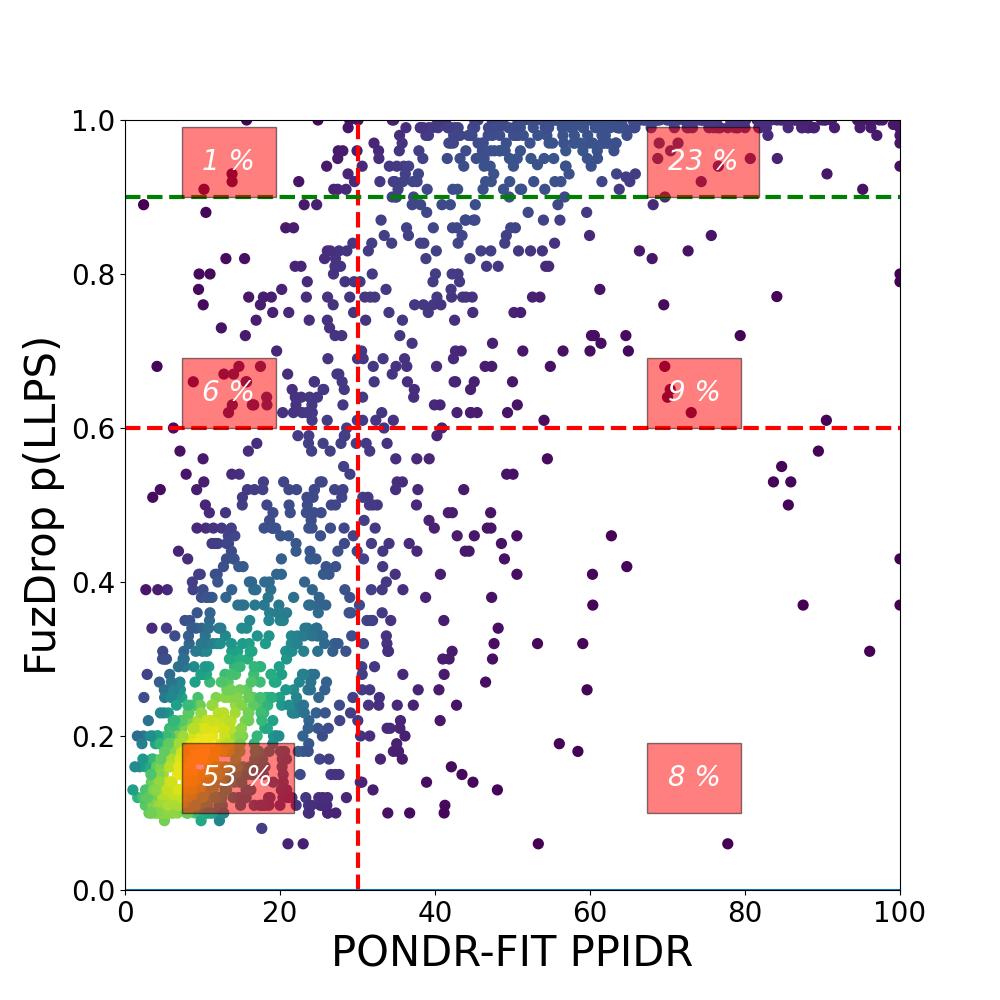 |
|  | Aging dataset | Control dataset |
|  | 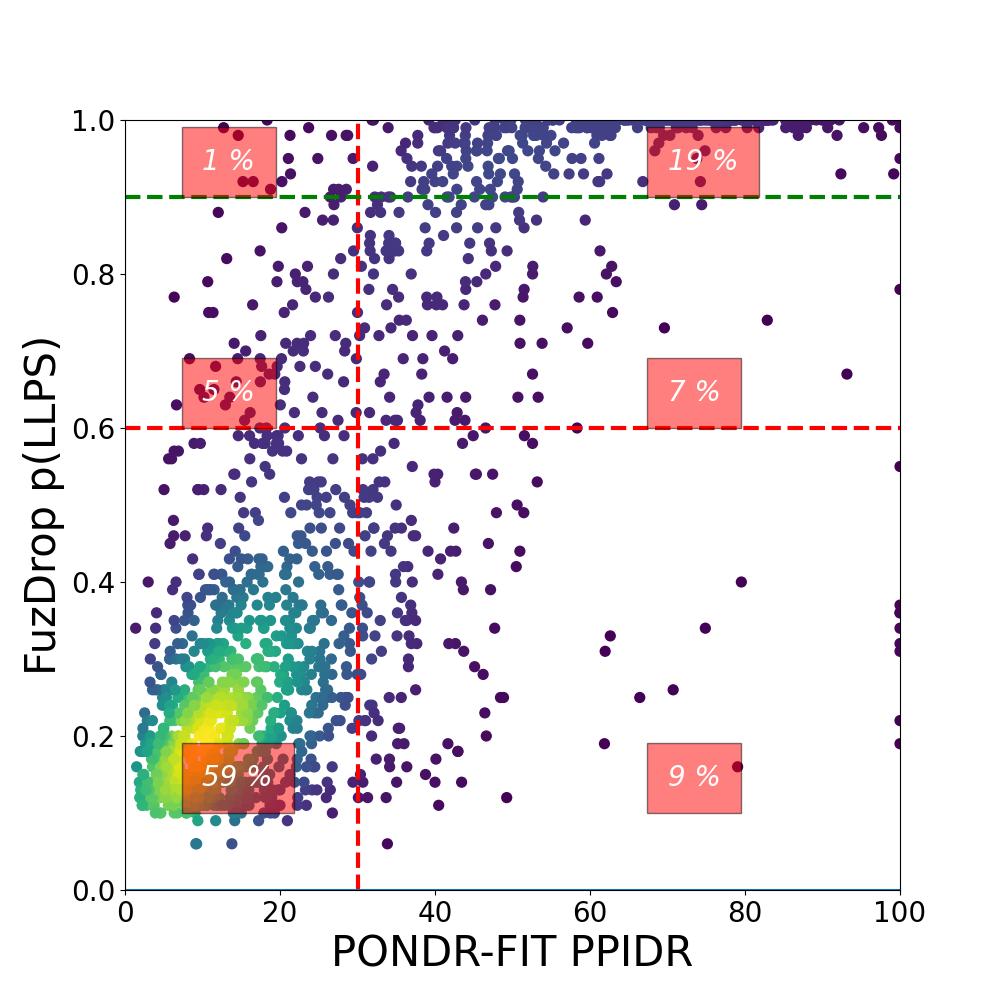 | 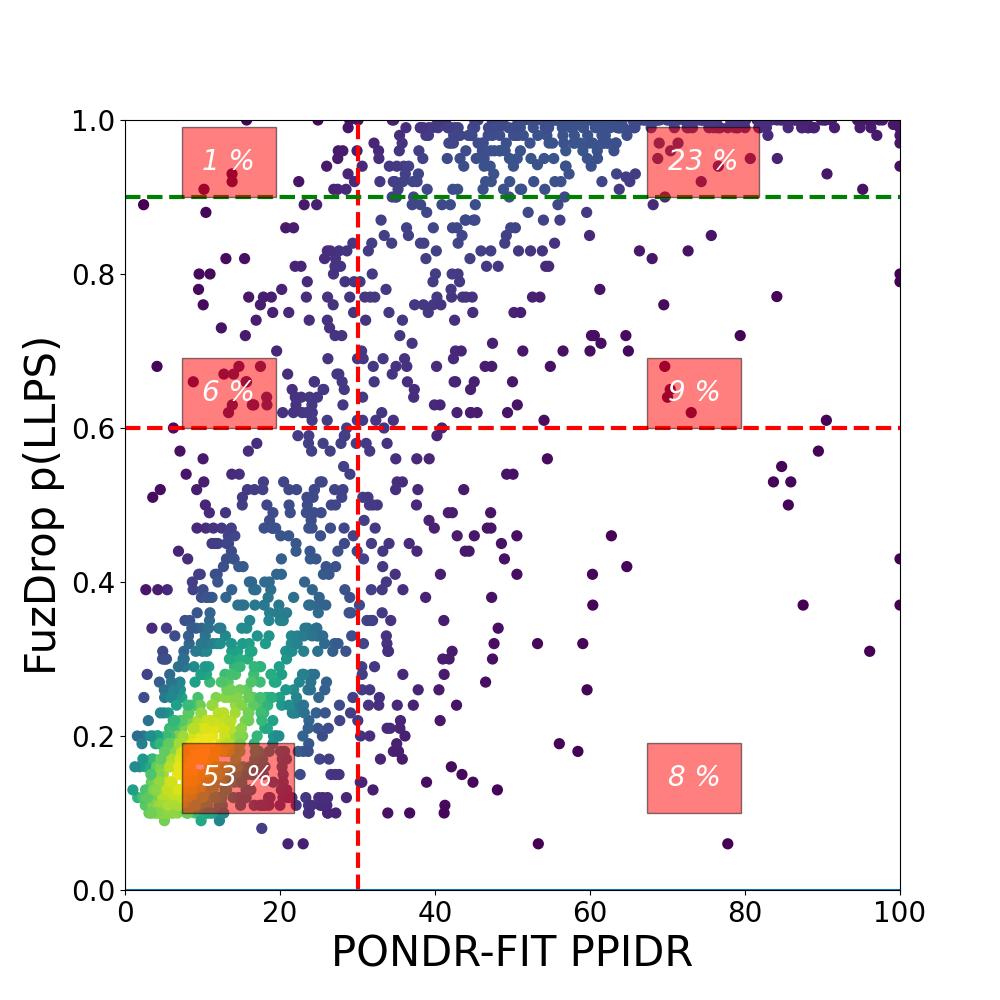 |

##### Figure S2. Disorder score correlates with MoRFs and LLPS propensity for the aging dataset.

##### A. 3 heat-maps for differently disordered groups (from left to right - IDPs, partial IDPs, ordered proteins). Proteins sorted by disorder score PPIDR^PONDR-FIT^ – highest on the top and smallest on the bottom, In each heatmap disorder state shown by constantly colored columns - red (for IDPs, 572 proteins), violet (for partial IDPs, 689 proteins), green (for ordered proteins, 363 ones) – placed left and right from column with PPIDR^PONDR-FIT^ values, which is gradually colored from red to green. On the left edge Percentage of MoRFs length (% of protein length) shown, on the right edge – two columns with p LLPS from PSPredictor and from FuzDrop. Trends of MoRFs, PPIDR^PONDR-FIT^ and p LLPS by PSPredictor and FuzDrop shown on Supplement Figure 1.

B. Values for 1624 proteins of aging dataset of MoRFs number, disorder score (PPIDR^PONDR-FIT^), LLPS propensity (by PSPredictor and FuzDrop). Proteins sorted by disorder score PPIDR^PONDR-FIT^.

C. Correlation matrix between PSP, FuzDrop score, MoRF, PPIDR^PONDR-FIT^. Pearson correlation R was used, which is correct because data has multivariable normal distribution.

##### D. Correlation between ID and LLPS propensity (FuzDrop)

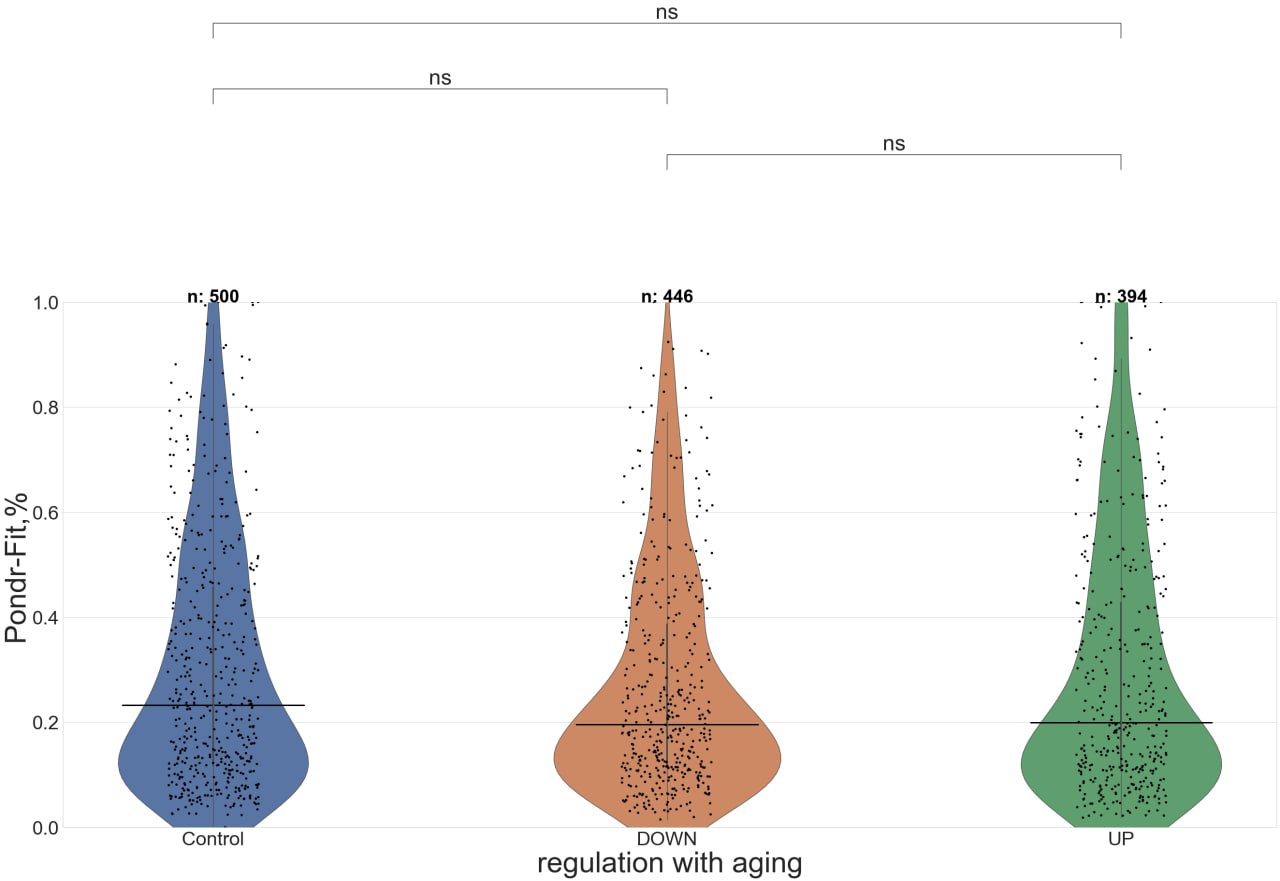

Figure S3. Proteins of the resulting dataset, for which the direction of expression change with aging progression is known, using information from literature [81, 82, 119]. UP-regulated and DOWN-regulated proteins did not show statistical differences in levels of intrinsic disorder from each other and the control sample (dataset of same size and random proteins). ns - not significant.

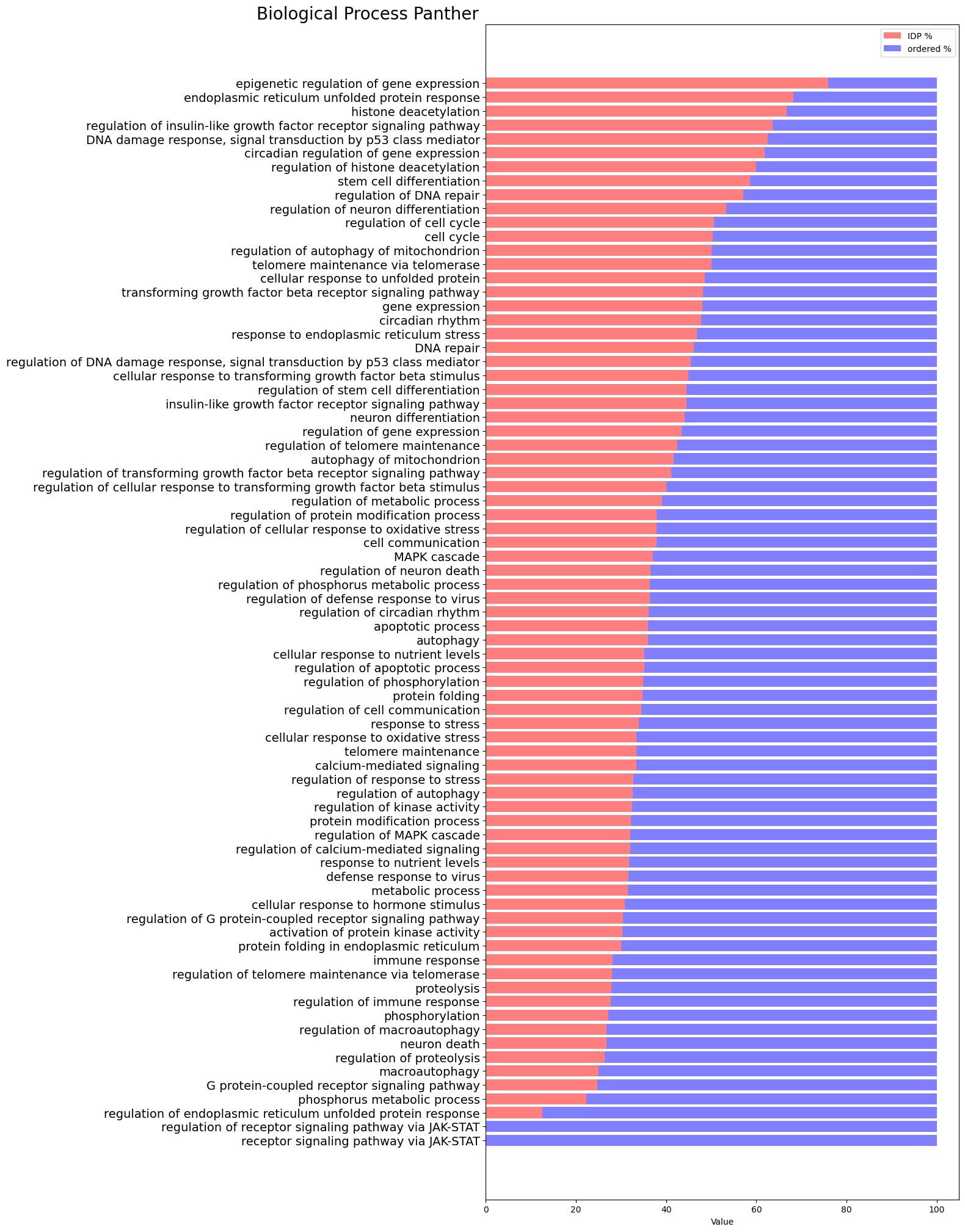

Figure S4. Analysis of presence of IDP in different processes by CH-CDF analysis. Ordered proteins assignment was done for ones having positive CDF and negative CH values (Q1 quartile in CH-CDF plot).

The ‘regulation of gene expression’ process is enriched in IDPs, as comparison of ageing-related and the rest proteins that is assigned to this process by Gene Ontology showed that there is 10% increase in the PPIDR median value of proteins associated with aging, compared with other proteins of this process (distribution of differences is normal, which is confirmed by the Shapiro-Wilk test, the confidence interval is (1.3; 18.9) (Figure S3A)). Bootstrapping was used to quantify the difference in median values in the distributions. It allows to determine the confidence interval, where difference between medians of the compared samples is located.

| A | B | C |
| --- | --- | --- |
| 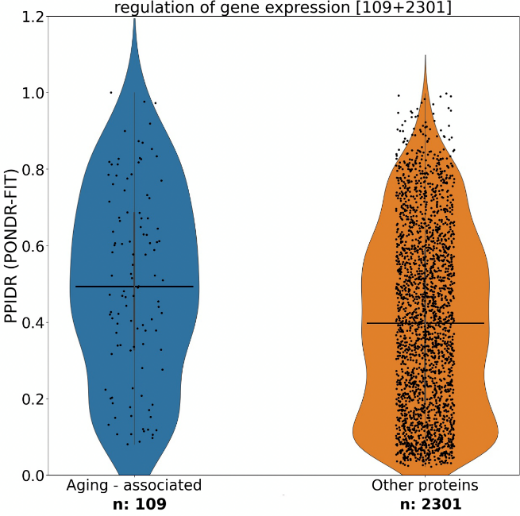 | 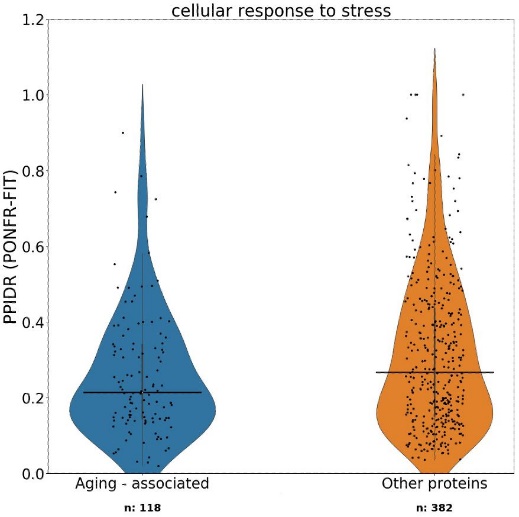 | 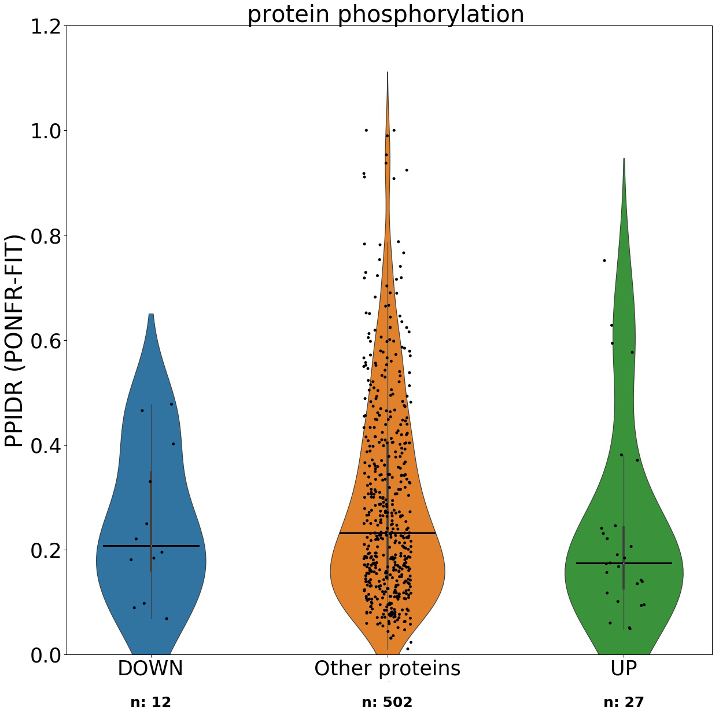 |
| 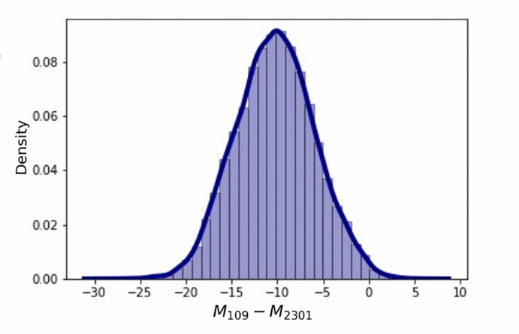 | 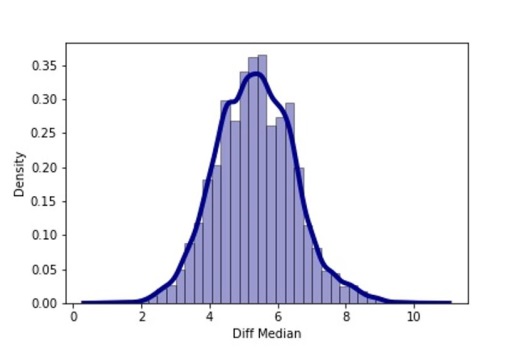 | 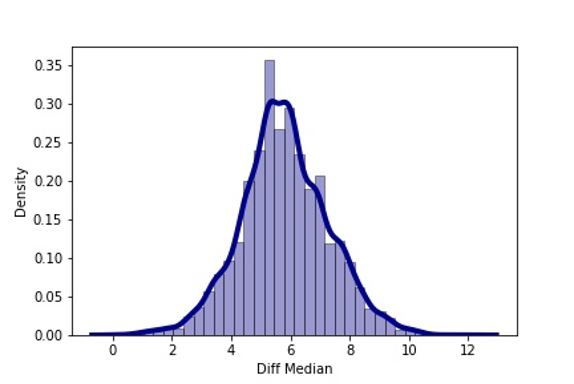 |

Figure S5. Comparison of aging-related proteins with other proteins in one BP on the median level of disorder.

A. Regulation of gene expression BP has higher disorder for aging-related proteins (in comparison with all other BP proteins).
B. Cellular response to stress BP has higher disorder for aging-related proteins (in comparison with all other BP proteins).
C. Protein phosphorylation BP has higher disorder for aging-related proteins (in comparison with all other BP proteins).

This analysis showed that for the ‘cellular stress response’ and ‘post-translational modification’ processes, statistically significant differences also are observed (Figure S3B, C). However, the median PPIDR of the proteins associated with aging for stress response and protein phosphorylation is approximately 5% lower. These results show that intrinsic disorder can be not only higher for ageing-related proteins, but also lower. **Therefore, the presence and absence of protein intrinsic disorder may be important for aging-related protein subgroup within the biological process.**

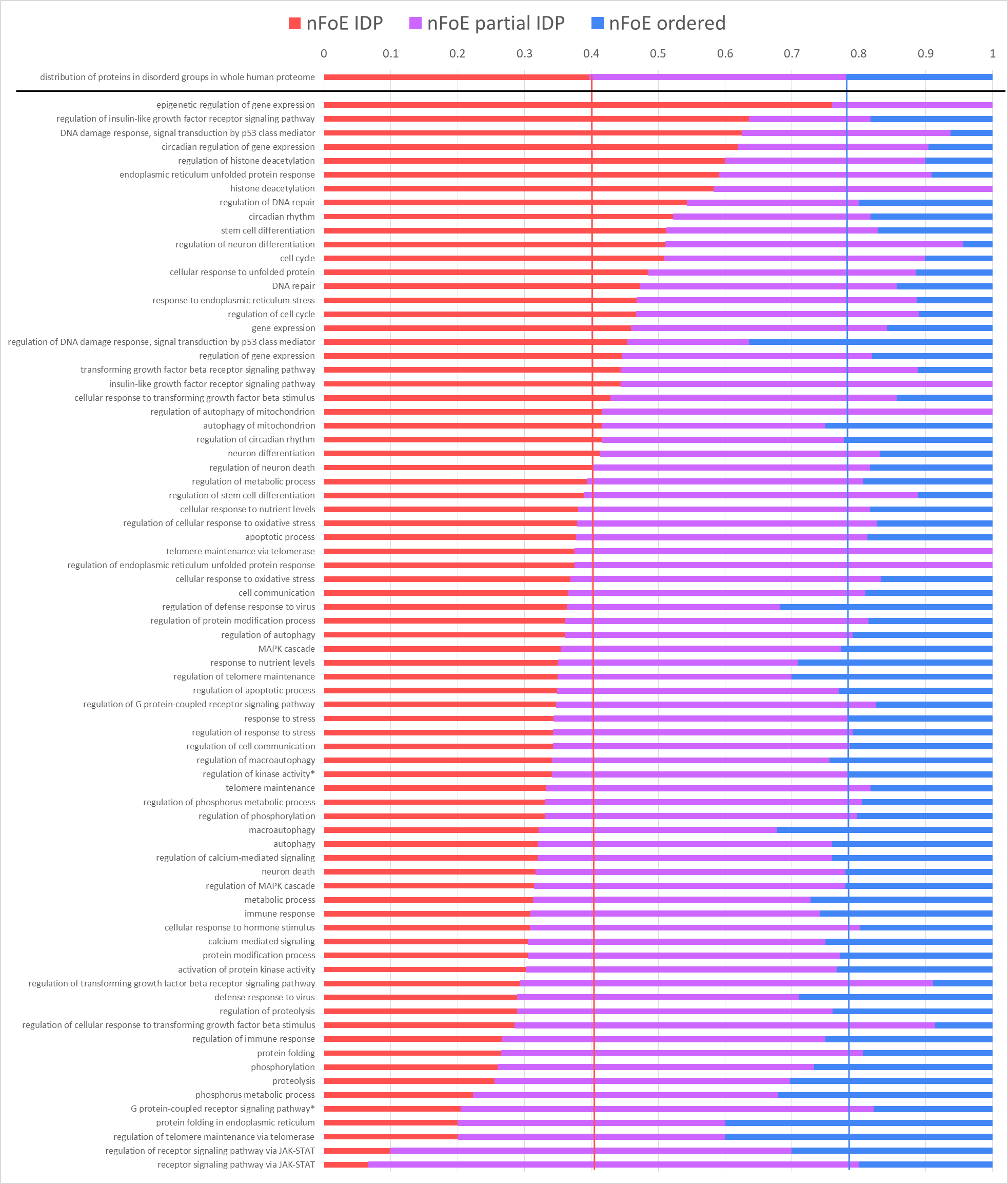

Figure S6. Distribution of aging-related proteins to BP with disorder assignment.

| Panther | BinGO |
| --- | --- |
| 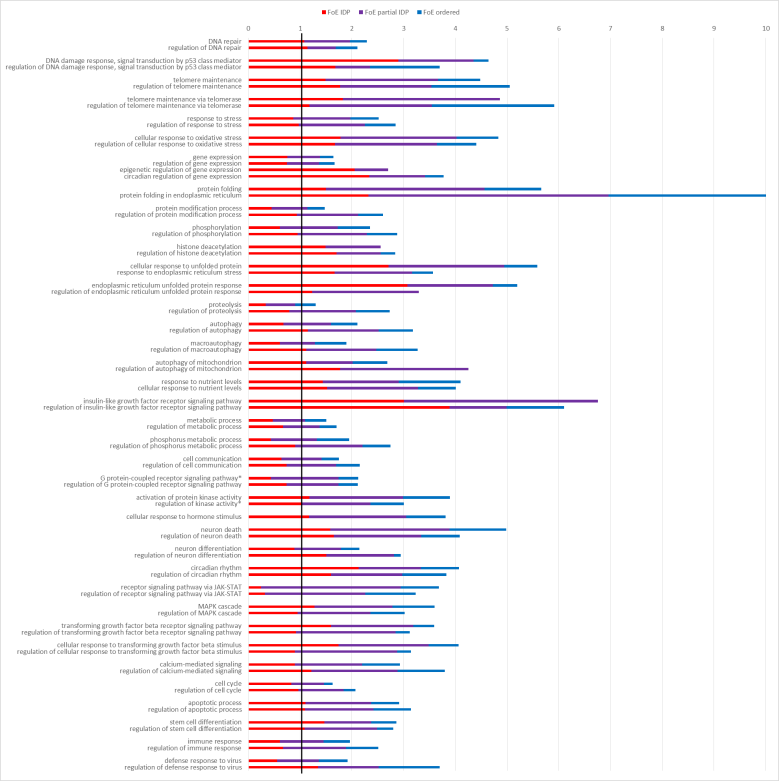 | 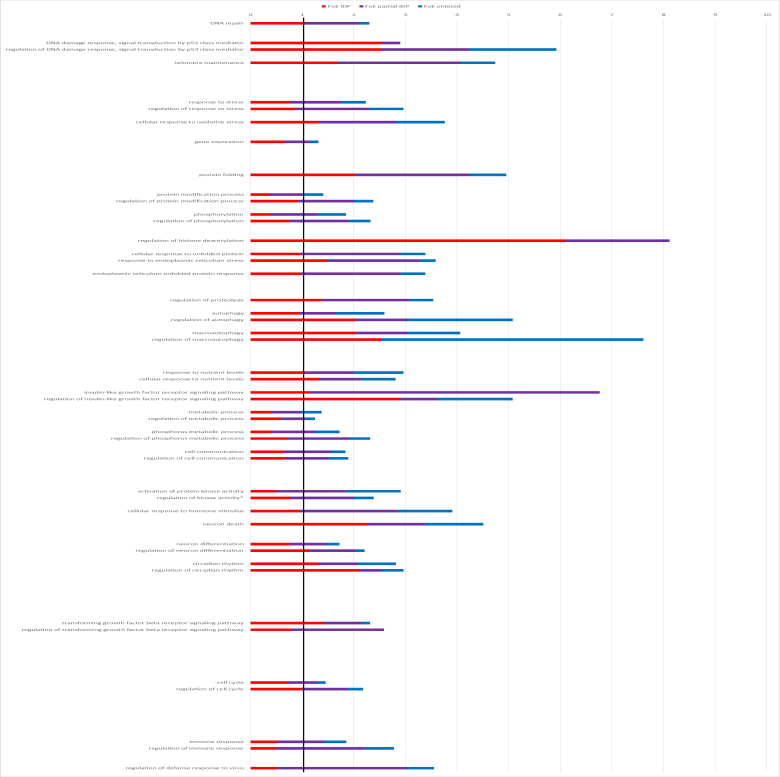 |
| Panther | BinGO |
| 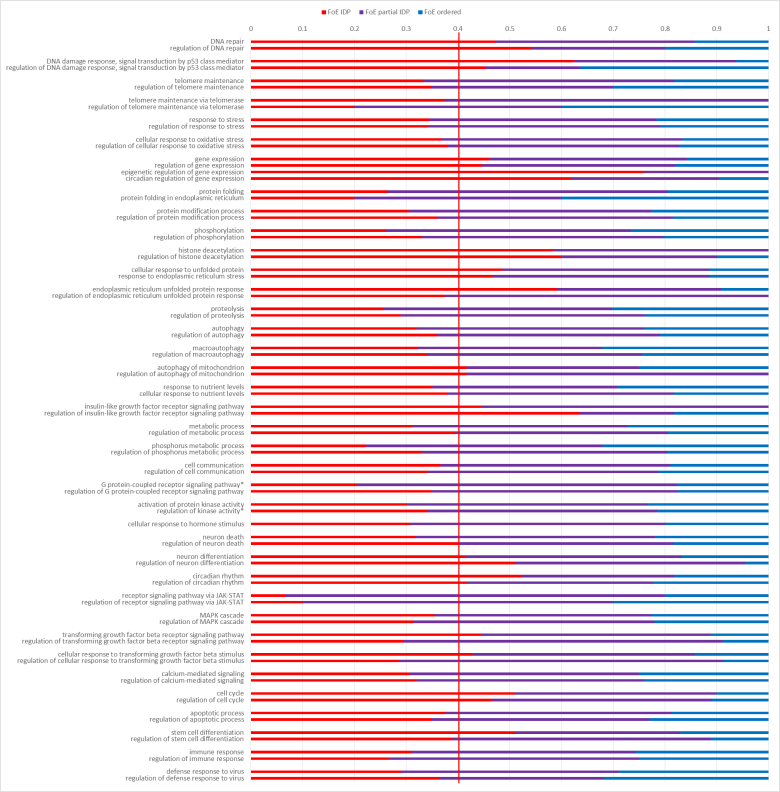 | 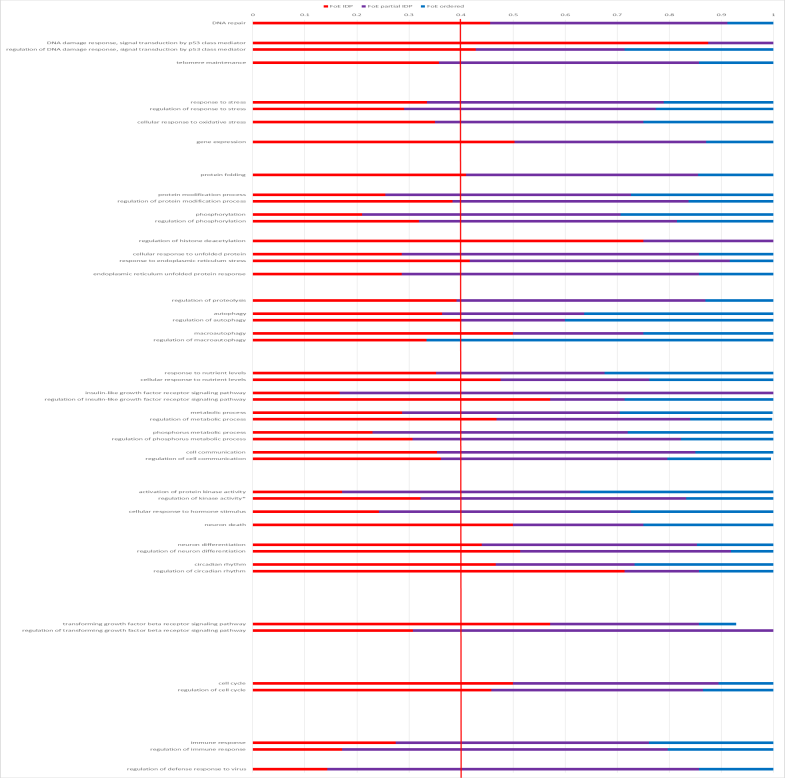 |

Figure S7. Comparison of enrichments (number of proteins in BP of interest for the Aging Proteome divided by expected number of proteins in BP of interest for the random dataset of the equal size as Aging Proteome) for different Biological Processes (BP), obtained from PantherDB and BinGO CytoScape App v3.0.5. All 1624 proteins from Aging proteome were analysed on GO Biological Processes for Homo Sapiens organism, overrepresentation, reference dataset was whole human proteome. Statistical hypergeometric test with Benjamini & Hochberg False Discovery Rate (FDR) correction was used with significance level 0.05.

Top panels represent not normalized enrichments, bottom ones are normalized.

Impact of different disordered classes to enrichment represented by coloring: blue – ordered, violet – partially disordered, red - IDP.

nFoE – normalized fold enrichment (number of proteins in BP of interest for the Aging Proteome divided by expected number of proteins in BP of interest for the random dataset of the equal size as Aging Proteome). If one will investigate highly disordered proteins, related to aging, amount of proteins involved in ‘DNA repair’ will be about 2 times higher, than expected for a random sample.

| A Control dataset 1  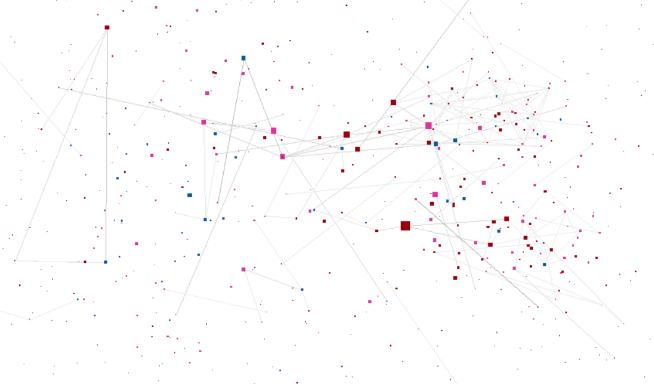  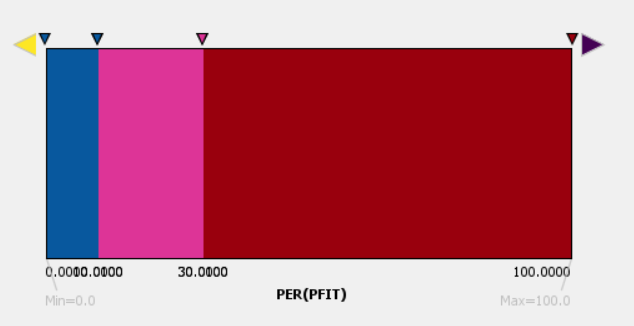 | |
| --- | --- |
| B. Aging dataset BinGO BP trees for clusters | |
| Cluster №1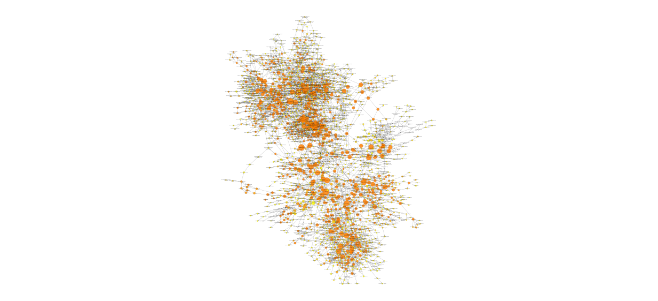 | Cluster №1-1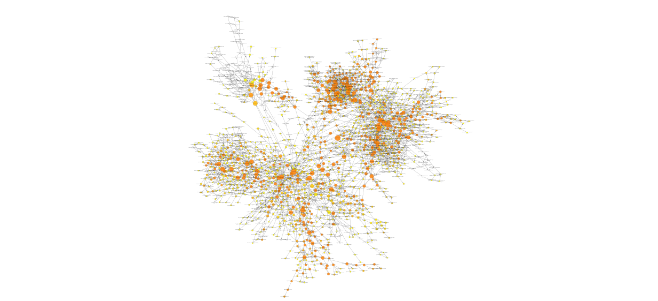 |
| Cluster №1-2  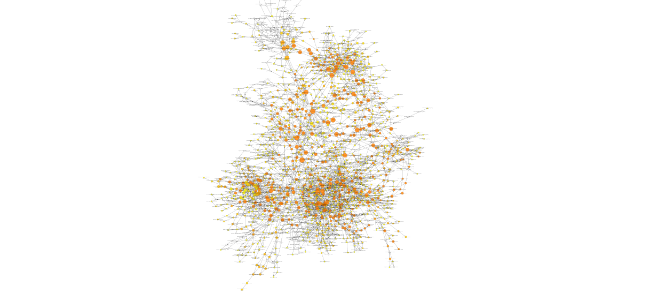 | Cluster №2 Down-regulated with aging 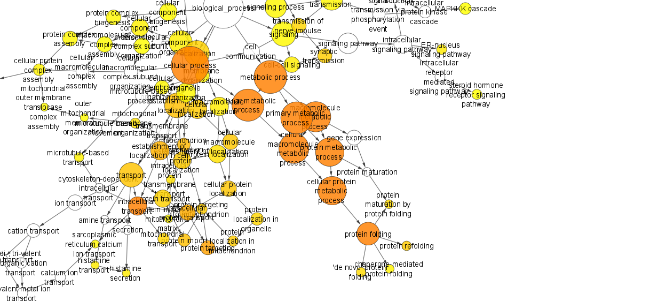 |
| Cluster №3  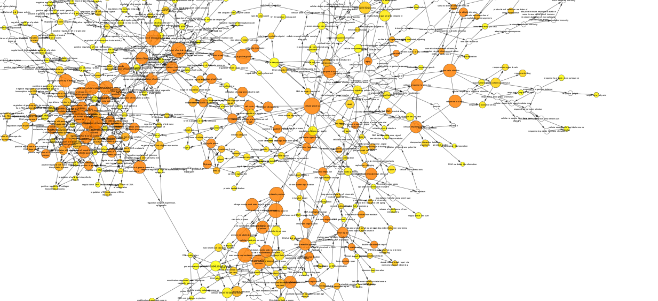 | Cluster №4 UP-regulated with aging  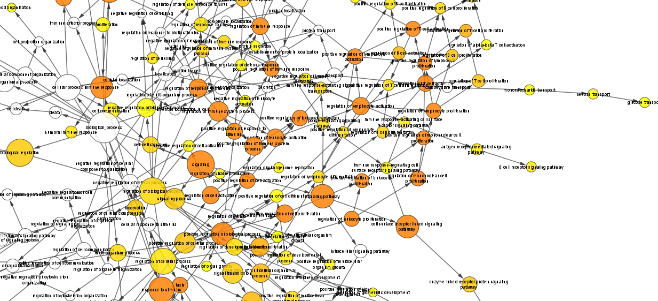 |

Figure S8. CytoScape results of proteins and Biological Processes (BP) analysis

A Protein-protein interaction network for control dataset of random 1624 proteins. Nodes colors reflect structural disorder: blue – ordered, violet – partially disordered, red - IDP. Size of nodes reflects their hubness (STRING degree value).

B. Bingo Results of GO BP overrepresentation in Aging dataset. BP overrepresentation shown in the gradient of orange color (the most dense orange reflects the minimal p-value).

| Cluster №1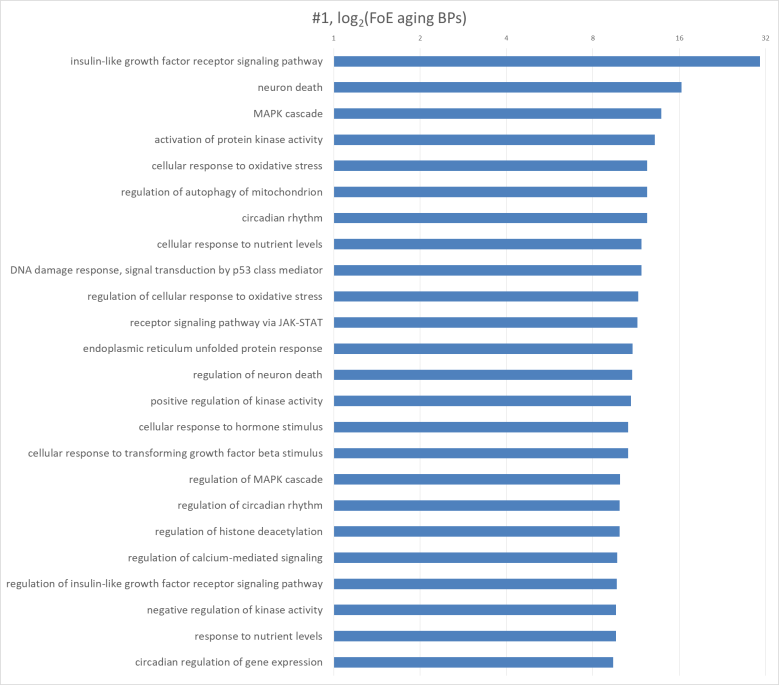 | Cluster №2 Down-regulated with aging  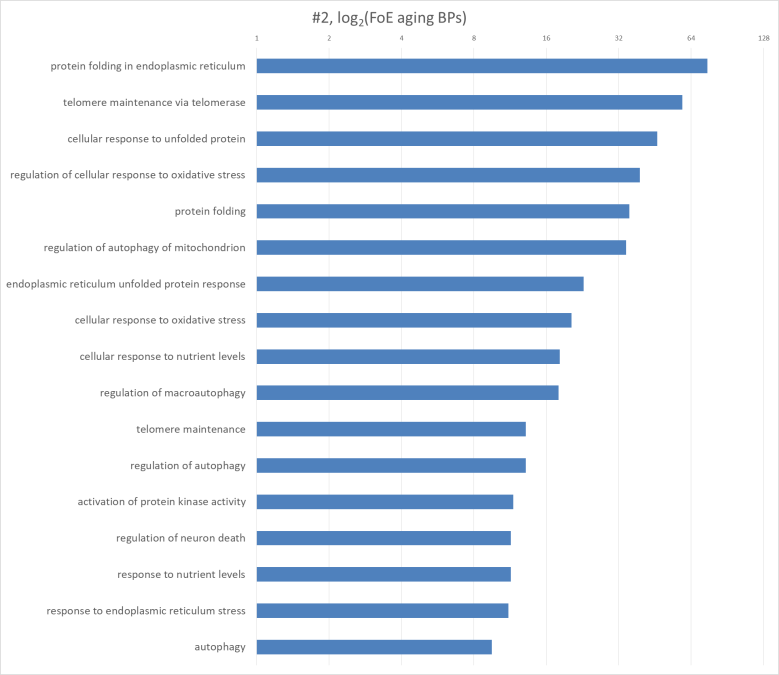 |
| --- | --- |
| Cluster №3 IDP, LLPS enriched  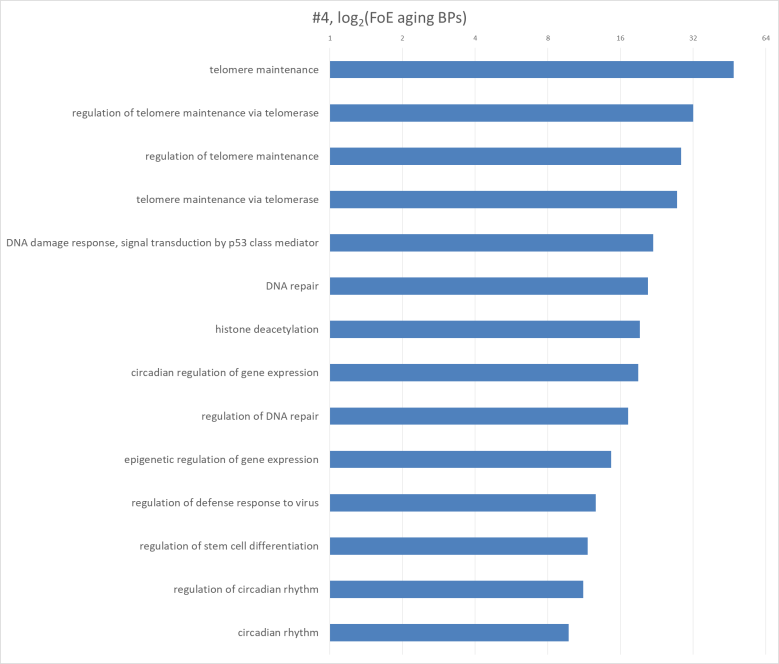 | Cluster №4 UP-regulated with aging  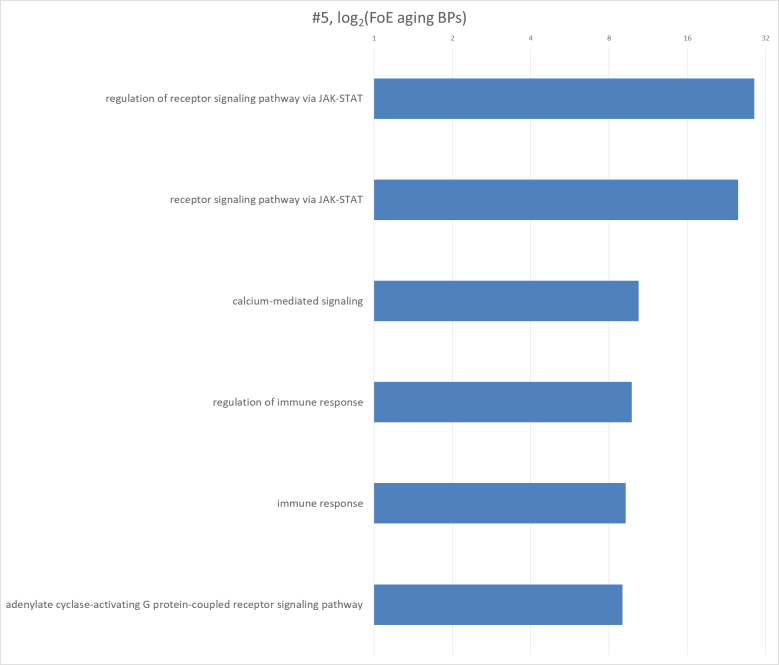 |

Figure S9. Panther Results of GO BP overrepresentation

| Cluster №1 | Cluster №2 Down-regulated with aging – protein folding  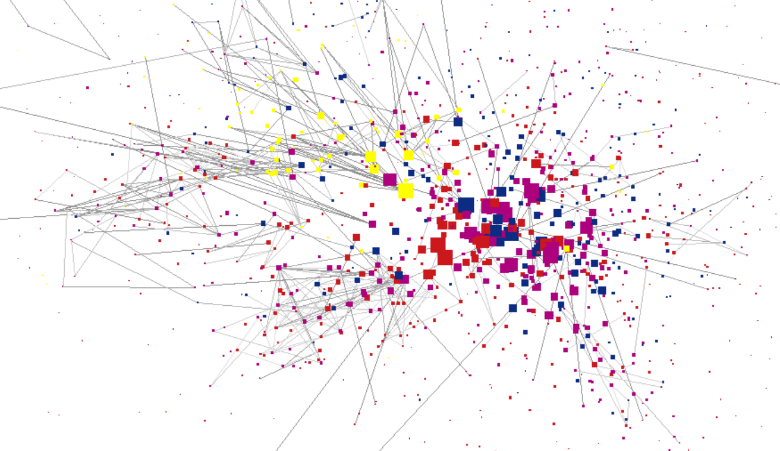  Down-regulated with aging  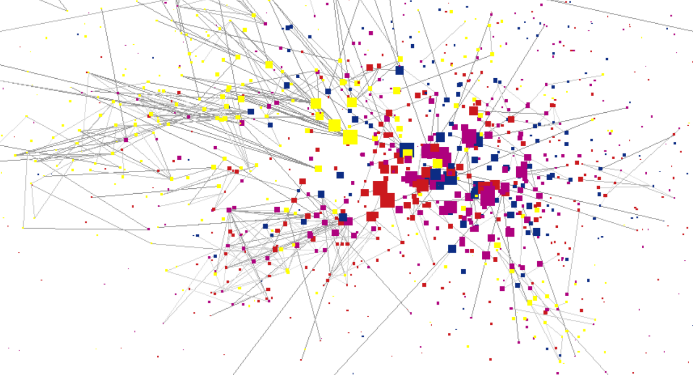 |
| --- | --- |
| Cluster №3 IDP, LLPS enriched  DNA metabolic process  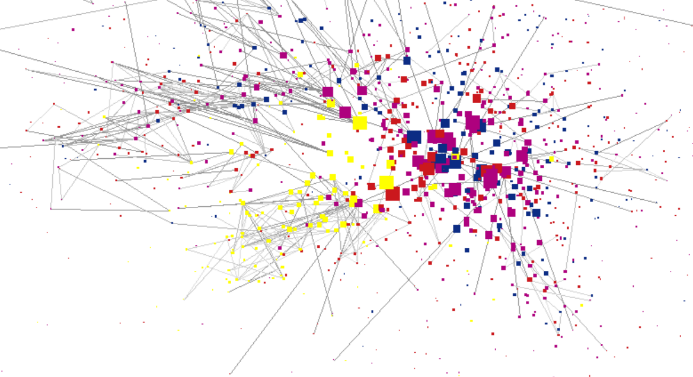  DNA repair  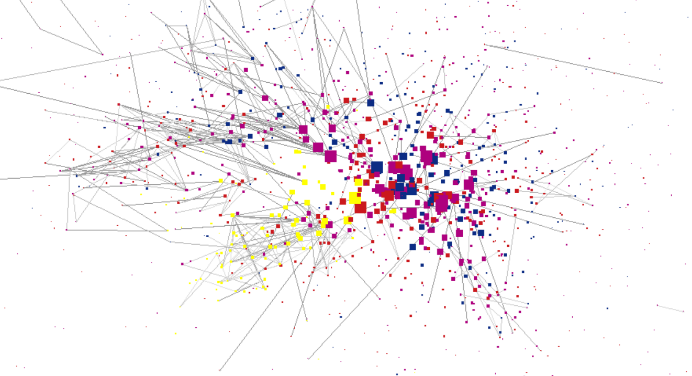  Cell cycle  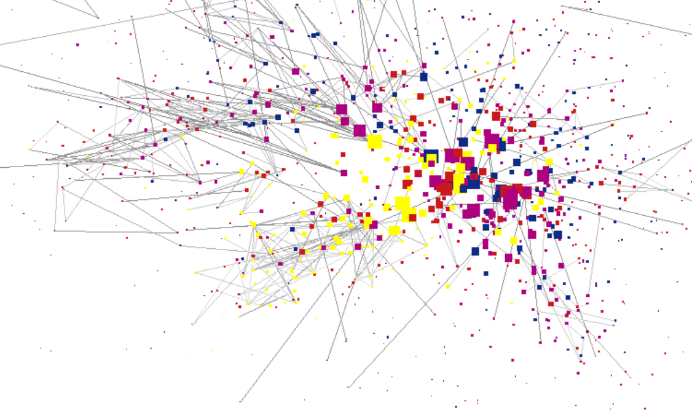  Telomere maintenance  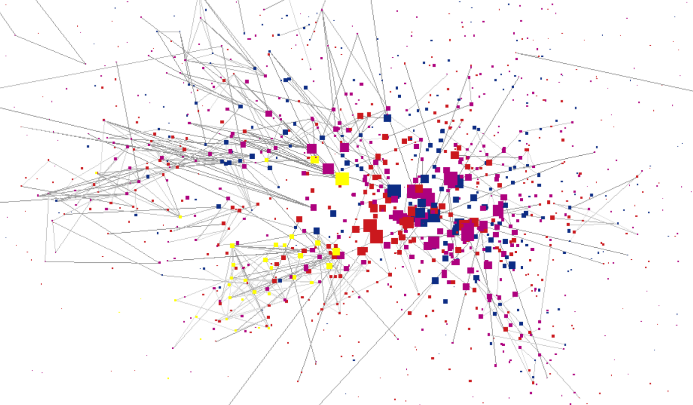 | Cluster №4 UP-regulated with aging  Adaptive immune response  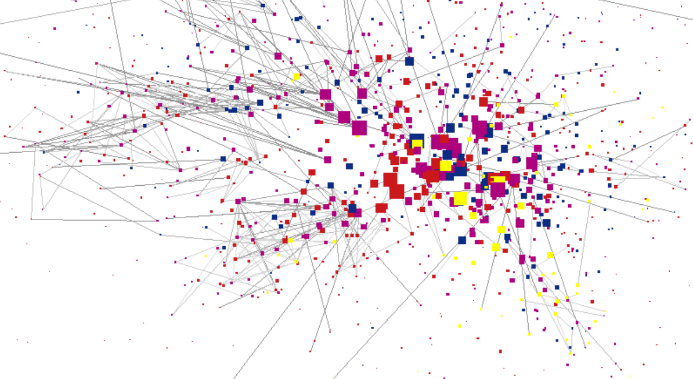  UP-regulated with aging  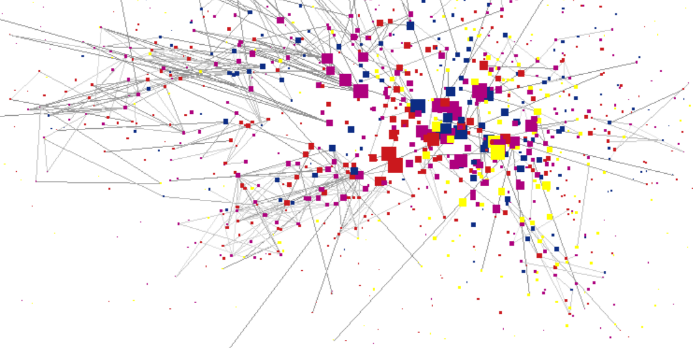 |

Figure S10. Panther Results of GO BP overrepresentation

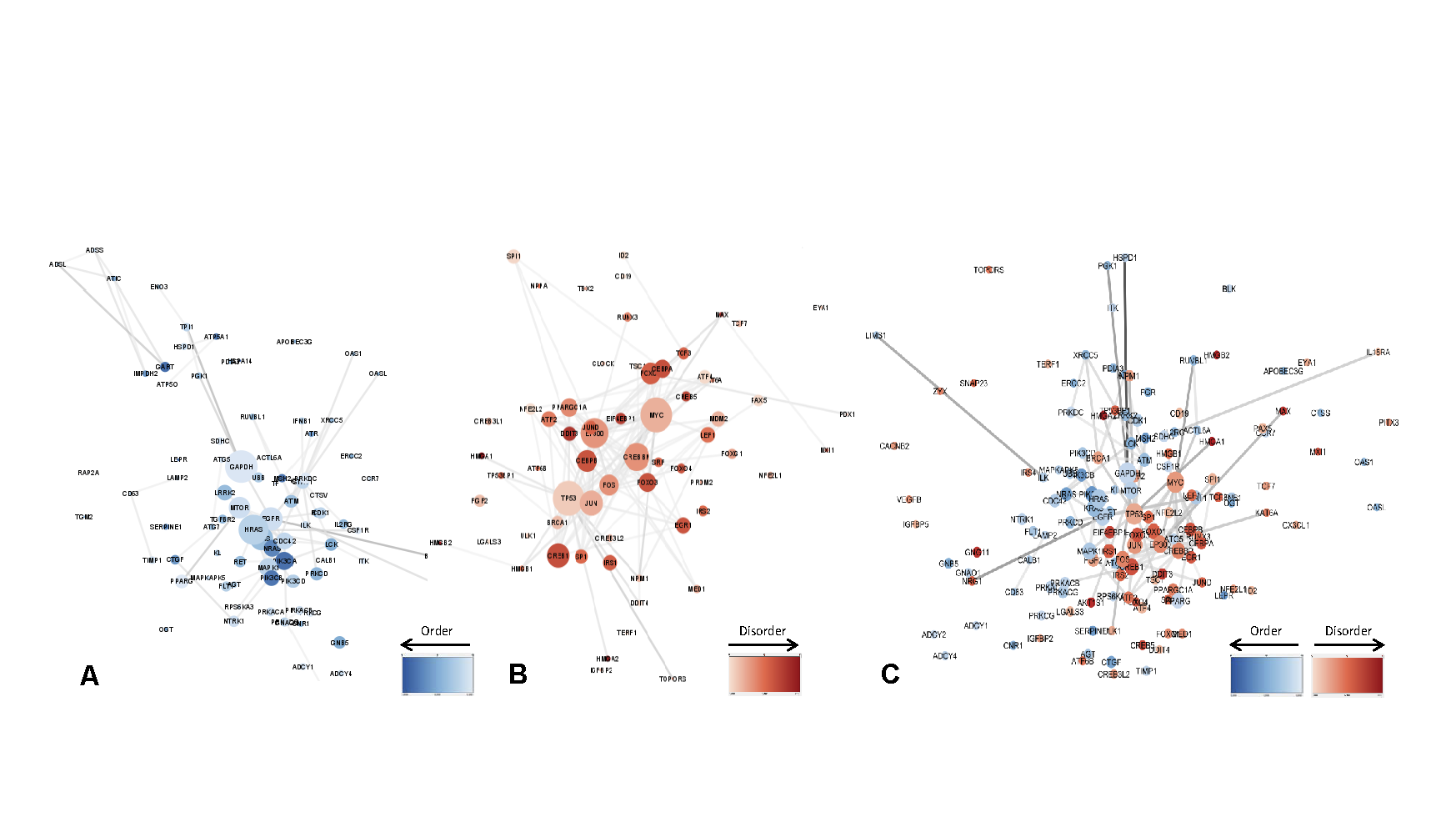

| ordered | disordered | disordered + ordered |
| --- | --- | --- |
| 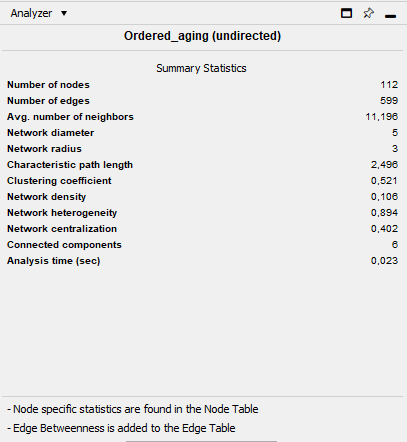 | 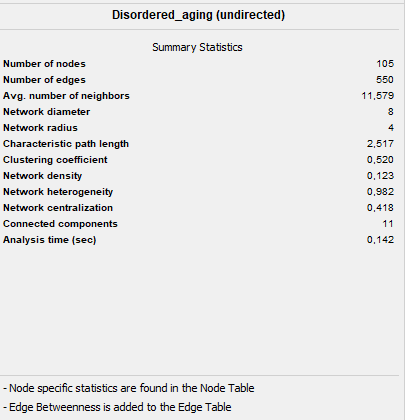 | 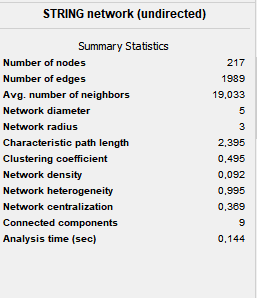 |

##### Figure S11. PPI-network of highly ordered (A), highly disordered (B) proteins and both of them (C).

### LLPS prediction in PPI network clusters

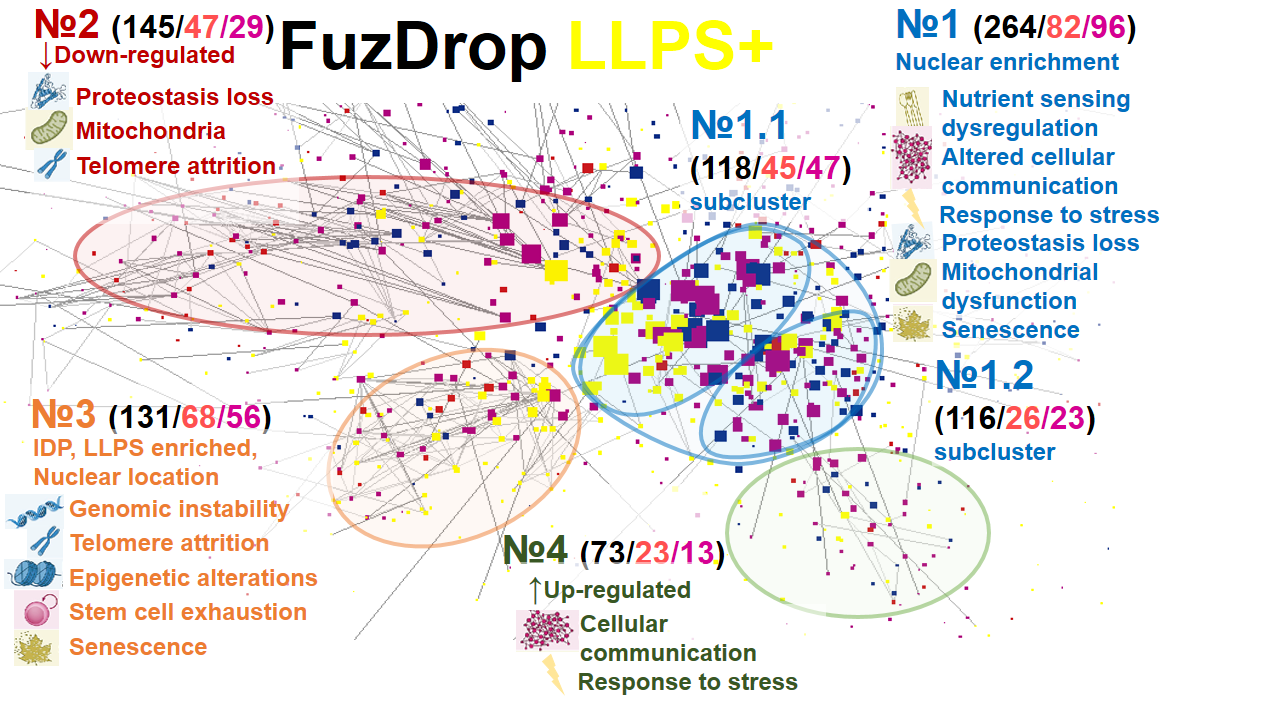

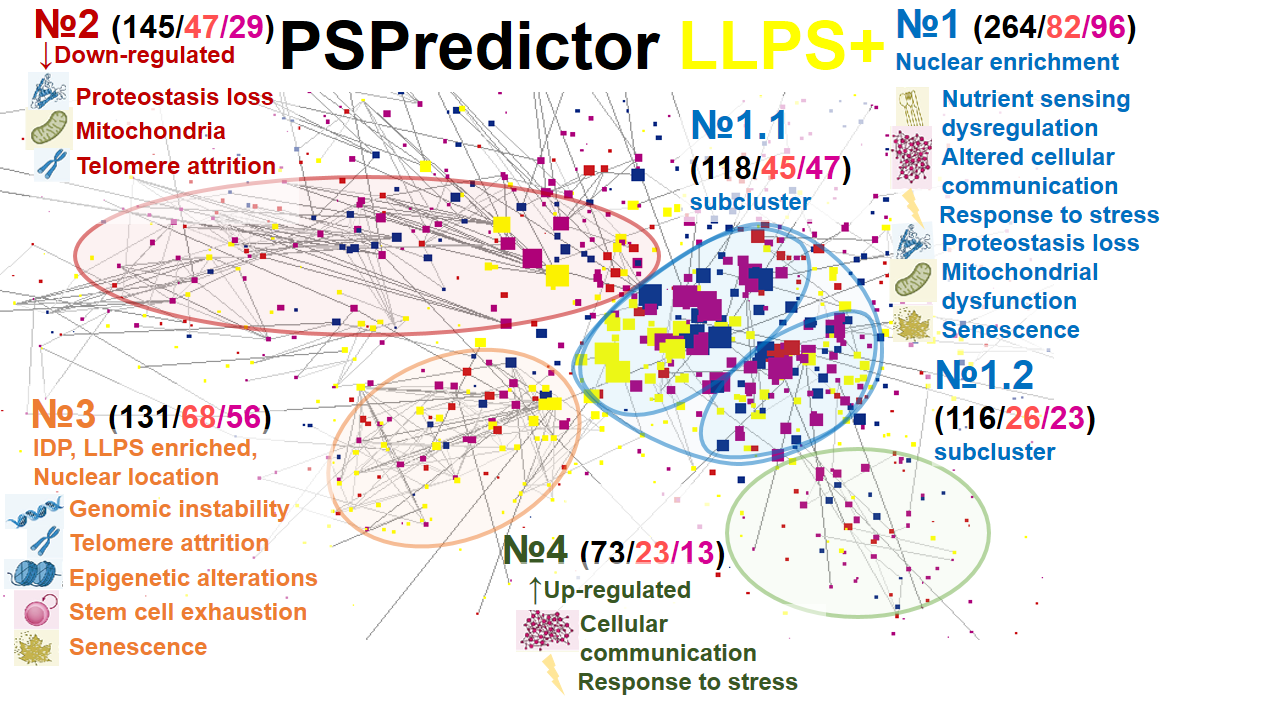

##### Figure S12. FuzDrop and PSPredictor results of LLPS prone proteins in CytoScape PPI network.

*Supplementary Note 1*

Interestingly, changes in plasma protein expression are often non-linear. In a study by Tony Wiss-Coray and colleagues (2019), the DE-SWAN bioinformatics program was developed. This algorithm analyzes protein levels over a period of 20 years and compares two groups of 10 years (eg, 35–45 years old versus 45–55 years old), shifting the window in 1-year increments from young to old. Hundreds of proteins have been identified using this algorithm, whose expression changes in waves with age. The summation of the number of differentially expressed proteins at each age revealed three peaks at ages 34, 60, and 78 years. The next challenge was to understand whether the proteins that make up the three waves of aging were uniquely linked to the genome or proteome of age-related diseases and traits. As it turned out, proteins associated with cardiovascular disease, Alzheimer's disease and Down's syndrome showed a strong overlap with proteins that make up the waves of aging [119].

#### Supplementary Note 2. Predictors for intrinsic disorder prediction

For the analysis of a large number of proteins, appropriate analytical methods are required. The following is a description of the set of computational algorithms utilized to analyze the intrinsic disorder in this study. This set contains algorithms that predict the level of disorder of the protein and the number of potential binding sites.

*Prediction of intrinsic disorder.*

1) PONDR VLXT is described in [137] and is available at http://www.pondr.com/. PONDR VLXT consists of three neural networks, two for the terminal ends of the amino acid sequence and one for the interior. The end result is the average over the results of these neural networks. The input data for neural networks are numerical structural characteristics for each amino acid residue [137]. PONDR-VLXT, being one of the first predictors of disorder, is characterized by high sensitivity to the features of the amino acid sequence in areas where there is an interaction of proteins, mediated by lack of structure [138]. Although this algorithm is not the most accurate, it is useful in identifying MoRFs, short, disordered regions capable of acquiring structure when interacting with partner proteins.
2) PONDR VL3 is described in [139] and is available at http://www.pondr.com/ and http://www.disprot.org/metapredictor.php. This algorithm uses a simple voting on several neural networks, which receive data on the composition, complexity and Shannon informational entropy of the input amino acid sequence [139]. PONDR VL3 works best on completely disordered proteins and for detecting long disordered regions.
3) PONDR VSL2 is described in [89, 140] and is available at http://www.pondr.com/. This algorithm was trained on 1,335 protein sequences containing 230 long disordered regions (25,958 residues), 983 short disordered regions (9,632 residues), and 354,169 ordered residues [89, 140]. PONDR VSL2 is based on the support vector machine. This algorithm receives data on the composition of the sequence, information on evolutionary conservatism, and a predicted secondary structure as input. PONDR VSL2 performs statistically better on proteins containing both ordered and disordered regions. For a long time, this program was one of the most accurate among autonomous analogues (Fan and Kurgan, 2014; Peng and Kurgan, 2012; Peng et al., 2005b).
4) The PONDR-FIT algorithm is described in [72]. PONDR-FIT integrates six individual algorithms (PONDR VLXT [137], PONDR VSL2 [140], PONDR VL3 [139], FoldIndex [141], IUPred [142], TopIDP [143]). It is more accurate than each of the algorithms separately [72]. Not statistically different from PONDR VL3 for fully disordered and fully structured proteins, this algorithm is slightly better (by 1 standard deviation) than PONDR VSL2 when both structure and disorder are present.
5) IUPred2A is available at http://iupred2a.elte.hu. This algorithm is described in [142, 144]. IUPred is a tool for predicting extended disordered regions from amino acid sequences by estimating their total pairwise interaction energy between residues. The algorithm is based on the assumption that disordered sequences do not acquire a clear structure due to their inability to form sufficient stabilizing interactions between residues [142, 144]. This tool can predict short or long regions of intrinsic disorder based on built-in optimized parameter sets [142, 144].
6) The ‘charge-hydropathy’ charting computer program is available at http://www.pondr.com/. The basic algorithm of this binary classifier is described in [50]. For the protein under consideration, the graph represents the dependence of its absolute average charge - excluding histidine - on the average hydropathy according to the Kyte-Doolittle scale [145]. The hydropathy score is on a scale of 0 to 1. The results of this analysis show that the elongated intrinsically disordered proteins are located in a unique region of phase space on the graph, with total charge and protein hydropathy (CH) as axes. This graph allows to separate ordered proteins and those without structure by a linear boundary.
7) Analysis of the cumulative distribution function (abbr. CDF) is another binary disorder classifier available at http://www.pondr.com/. This predictor is described in [73, 94]. In this analysis, the data for the protein in question are shown in the form of an CDF curve that plots PONDR values against their cumulative frequency for that sequence. This allows to discriminate between ordered and disordered proteins based on the distribution of values obtained from the algorithms. According to CDF analysis, fully disordered proteins have a very low percentage of residues with low predicted disorder scores. For ordered proteins, in contrast, most residues are assumed to have low disorder values. Therefore, in theory, all completely disordered proteins should remain in the lower right corner of the CDF plot, while all completely ordered proteins should be located in the upper left corner of this plot [73, 94].
8) CH-CDF analysis divides all proteins into four groups: structured, mixed, disordered and rare. It is based on the simultaneous use of data from the CH (Charge-Hydropathy) and CDF (Combined Distribution Function) approaches, which divide proteins into ordered and disordered [92]. The CH method was developed by Uversky et al. [50] based on the following model: repulsion from charges of the same sign promotes protein unfolding, while increased hydrophobicity promotes protein folding. The cumulative distribution function separates proteins into folded and unfolded proteins based on the distribution of predicted disorder values [146].

Additional value of this analysis comes from the fact that the two predictors consider intrinsic disorder from two different sides, complementing each other. The ‘CH’ classifier only uses information about total protein charge and its averaged hydropathy to separate proteins in two groups: those containing long disordered regions of disorder (native coils and native pre-molten globules) and proteins having compact globular conformation (native molten globules and ordered proteins). CDF classifier uses different approach: it separates ordered proteins from all proteins having any kind of intrinsic disorder (native globules, native molten globules and native pre-molten globules).

***Supplementary Notebooks***

Automated obtaining of the proteins UniProtIDs, Fold enrichments, p-values, GO terms from xml files form Panther DataBase was carried out using the Python libraries (see Supplement Notebooks).
